# Linking molecular pathways and large-scale computational modeling to assess candidate disease mechanisms and pharmacodynamics in Alzheimer’s disease

**DOI:** 10.1101/600205

**Authors:** Leon Stefanovski, Paul Triebkorn, Andreas Spiegler, Margarita-Arimatea Diaz-Cortes, Ana Solodkin, Viktor Jirsa, Anthony Randal McIntosh, Petra Ritter, for the Alzheimer’s Disease Neuroimaging Initiative

**Affiliations:** Brain Simulation Section, Department of Neurology, Charité - Universitätsmedizin Berlin & Berlin Institute of Health, Germany; Institute of Informatics, Free University Berlin, Germany; Anatomy & Neurobiology and Neurology, UC Irvine Health, Irvine, CA, USA; Institut de Neurosciences des Systèmes, Aix Marseille Université, Marseille, France; Baycrest Health Sciences, Rotman Research Institute, Toronto, Ontario, Canada; Bernstein Center for Computational Neuroscience Berlin, Germany

**Keywords:** Alzheimer’s disease, neurodegenerative disease, The Virtual Brain, PET, beta amyloid, EEG, MRI, excitotoxicity, NMDA, memantine, disease mechanisms, drug targets, personalized medicine

## Abstract

**Introduction:** While the prevalence of neurodegenerative diseases associated with dementia such as Alzheimer’s disease (AD) increases, our knowledge on the underlying mechanisms, outcome predictors, or therapeutic targets is limited. In this work, we demonstrate how computational multi-scale brain modelling links phenomena of different scales and therefore identifies potential disease mechanisms leading the way to improved diagnostics and treatment.

**Methods:** The Virtual Brain (TVB; thevirtualbrain.org) neuroinformatics platform allows standardized large-scale structural connectivity-based simulations of whole brain dynamics. We provide proof of concept for a novel approach that quantitatively links the effects of altered molecular pathways onto neuronal population dynamics. As a novelty, we connect chemical compounds measured with positron emission tomography (PET) with neural function in TVB addressing the phenomenon of hyperexcitability in AD related to the protein amyloid beta (Abeta). We construct personalized virtual brains based on individual PET derived distributions of Abeta in patients with mild cognitive impairment (MCI, N=8) and Alzheimer’s Disease (AD, N=10) and in age-matched healthy controls (HC, N=15) using data from ADNI-3 data base (http://adni.lni.usc.edu). In the personalized virtual brains, individual Abeta burden modulates regional inhibition, leading to disinhibition and hyperexcitation with high Abeta loads. We analyze simulated regional neural activity and electroencephalograms (EEG).

**Results:** Known empirical alterations of EEG in patients with AD compared to HCs were reproduced by simulations. The virtual AD group showed slower frequencies in simulated local field potentials and EEG compared to MCI and HC groups. The heterogeneity of the Abeta load is crucial for the virtual EEG slowing which is absent for control models with homogeneous Abeta distributions. Slowing phenomena primarily affect the network hubs, independent of the spatial distribution of Abeta. Modeling the N-methyl-D-aspartate (NMDA) receptor antagonism of memantine in local population models, reveals potential functional reversibility of the observed large-scale alterations (reflected by EEG slowing) in virtual AD brains.

**Discussion:** We demonstrate how TVB enables the simulation of systems effects caused by pathogenetic molecular candidate mechanisms in human virtual brains.

## 1. Introduction

Neurodegenerative diseases (NDD) gain increasing socioeconomic relevance due to an ageing society (1–4). The Alzheimer’s Association’s latest report estimates the yearly cost of Alzheimer’s disease (AD) treatment in the U.S. at $277 billion (5). By 2050 this number is expected to rise as high as $1.1 trillion. According to the report, early diagnosis could save up to $7.9 trillion in cumulated medical and care costs by the year 2050. While the prevalence of AD - the most common cause of dementia and the most common NDD in general - increases, its cause is still not understood, nor is there a cure. Our understanding of their pathogenesis and classification remain insufficient. Therefore, we aim to integrate clinical data from molecular biology and neurology, using nonlinear systems theory. Our aim is to build predictive models for health-outcome and cognitive function by individual virtual brain simulations using The Virtual Brain (TVB; thevirtualbrain.org) platform (6, 7). TVB integrates various empirical data in computational models of the brain that allow for the identification of neurobiological processes that are more directly linked to the causal disease mechanisms than the measured empirical data. Biomedical sciences are currently lacking a mapping between the degree and facets of cognitive impairments, biomarkers from high-throughput technologies, and the underlying causal origins of NDD like AD. The imperative for the field is to identify the features of brain network function in NDD that predict whether a person will develop dementia. The heterogeneity of NDD makes it difficult to develop robust predictions of cognitive decline. This can be addressed by large prospective studies where there is potential for participants to develop NDD. It is difficult in general to predict individual disease progression and this is a particular challenge in complex nonlinear systems, like the brain, where emergent features at one level of organization (e.g., cognitive function) can come about through the complex interaction of subordinate features (e.g., network dynamics, molecular pathways, gene expression). The Virtual Brain takes into account the principles of complex adaptive systems and hence poses a promising tool for identifying mechanistic predictive biomarkers for NDD. Due to the high dimensionality of brain models and the even greater complexity of the to-be-simulated brain states, selecting the used modeling approach carefully for a specific question of interest is essential.

The candidate biological mechanism under investigation in the present study is related to amyloid beta (Abeta), a protein that is an oligomeric cleavage product of the physiological amyloid precursor protein (APP) (8, 9). The soluble oligomers have the tendency for polymerization (9, 10). Due to their non-physiological configuration they aggregate and accumulates in brain tissue - a process that starts already in early preclinical stages of AD, i.e. many years before the onset of symptoms - typically in the fifth decade of life (11) - as shown in rodent models (12) and human studies (13, 14). Aggregated Abeta and its intermediates, soluble Abeta oligomers, can act directly neurotoxic (9, 15, 16) and have been found intra- or extracellularly (9, 15, 17). Those findings led to the hypothesis that the deposition of Abeta poses an initial step in the pathology of AD while Abeta has been suggested as a key feature in the pathogenesis of AD leading to major changes in the functionality and structure of the brain (13, 14, 18). The goal of the present study is to incorporate the hypothesized qualitative and quantitative effects of Abeta on neuronal population dynamics into our brain network models, i.e. adding mathematical models that describe how molecular changes alter population activity – so called cause-and-effect models. We will focus here on the disrupted inhibitory function of interneurons and consecutive hyperexcitability caused by Abeta – while we are aware of various other factors with potential roles for AD aetiology, such as vascular changes (19–21), neuroinflammation (22–25), genetics (26–28), environmental factors (29, 30) and concomitant proteinopathies others than Abeta pathology (31, 32). Beside Abeta there is a second molecular hallmark associated with the pathogenesis of AD: the phosphorylated Tau ‘tubulin-associated unit’ protein (8, 33, 34) which contributes to microtubule stability in the neural cytoskeleton (34). One major argument in favor of the more prominent involvement of Abeta in the pathogenesis of AD, in contrast to Tau, is its higher specificity to AD and its appearance in the early familial variants of AD, where the molecular pathway is better understood (14, 18, 35). Therefore, most therapeutic strategies in the past targeted Abeta. Yet recently three clinical trials with antibodies against Abeta had to be terminated in phase III: aducanumab (36, 37), crenezumab (38, 39) and solanezumab (40, 41) did not meet the expectations to act in a disease-modifying manner slowing down the cognitive decline (9). Nevertheless, there are still studies ongoing, e.g. with BAN-2401 (42). A relevant percentage of clinically diagnosed AD patients show additional brain pathologies beside Abeta and Tau in autopsy (32). Even in the cases of neuropathological AD diagnosis (i.e. secured Abeta and Tau pathology in histology), 55% of cases also exhibited a pathology of alpha synuclein (which we would expect in synucleinopathies like Parkinson’s disease) and 40% showed transactive response DNA binding protein 43kDa (TDP-43), a protein which we would expect in frontotemporal dementia or amyotrophic lateral sclerosis (31). Brain tissue of people who did not had relevant neurodegenerative brain changes in histological exams after death were showing Abeta in 50% and Tau pathology in 93% of the cases when using sensitive immunohistochemistry methods (31). Although Abeta and Tau are widely accepted as involved parts in the pathogenesis of AD and also define the disease entity (43), it remains unclear if they might be only epiphenomena of other contributing factors. This study hypothesizes a mechanistic role of Abeta in the disease process and builds a link between the molecular pathway alteration that leads to Abeta phenomenon of disinhibition and neural slowing in EEG **(Figure 1)**. Our mechanistic modeling approach can help to understand the complex inter-dependencies between the involved factors in AD and will improve through iterative refinement.

**Fig. 1.**
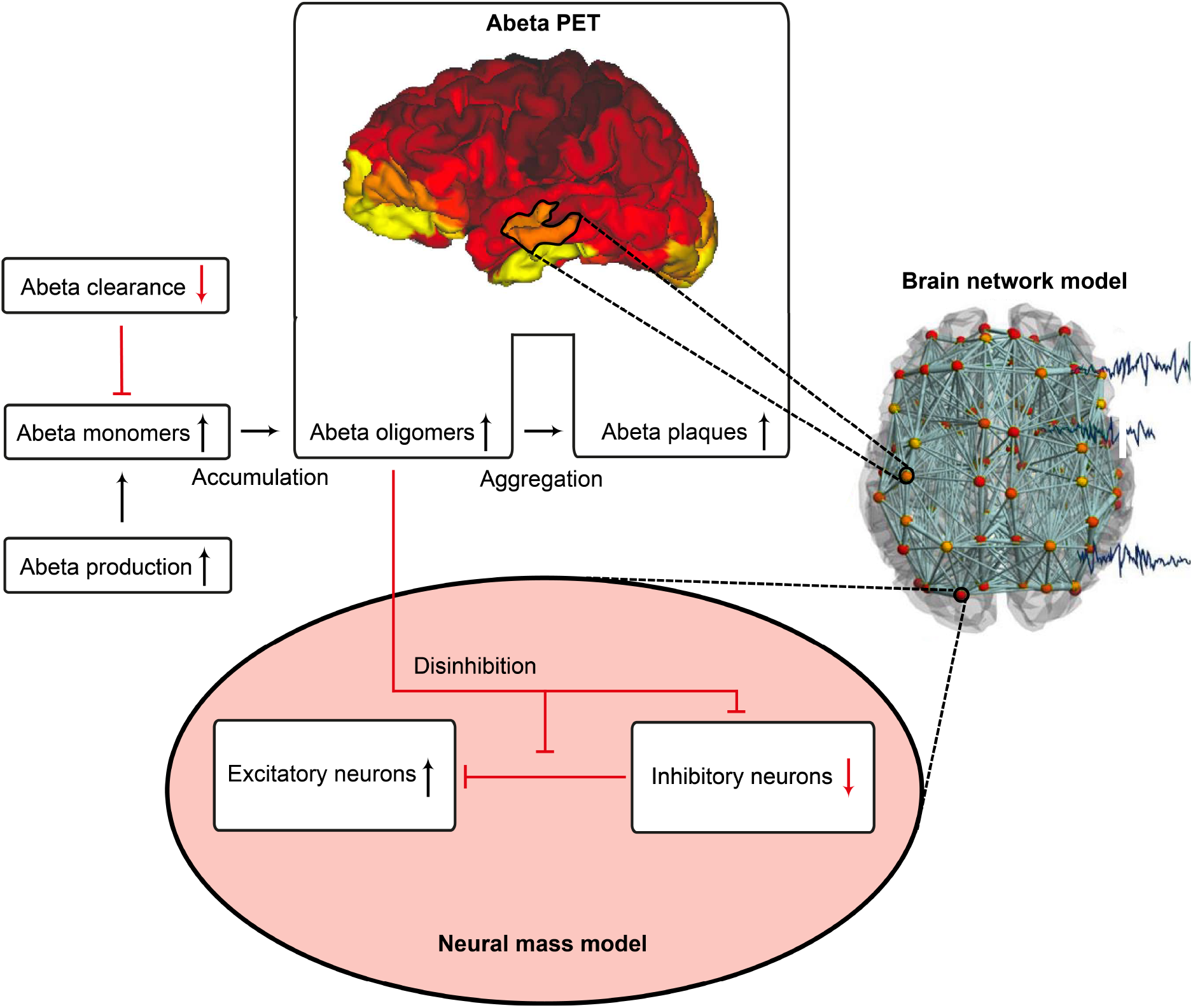
Cause and effect model: Alteration of the molecular Abeta pathway in AD cause disinhibition in the neural mass model. An altered pathway from soluble Abeta monomers to oligomers to insoluble plaques leads to potentially neurotoxic Abeta accumulation (9, 15, 16) that can be quantified by PET. Region specific Abeta burden leads to disinhibition in the neural mass model (44–48) - thus building a bridge between molecular pathways and brain network modelling. Parts of the figure are modified from (62).

Near Abeta plaques, a shift in neural activity has been observed (44). In AD mouse models with overexpression of APP and Presenilin-1, the number of hyperactive neurons was increased near Abeta plaques. This shift in the neuronal activity was associated with decreased performances in memory tests. Neuronal hyperactivity could be reduced by GABA agonists, suggesting pathology due to impaired inhibition. In neocortical and dentate gyri, pyramidal cells have been found to increase network excitability in vivo in an AD mouse model with overexpression of Abeta, that led to membrane depolarization and increased firing rates. A study by Hazra et al. (45) investigated an AD mouse model by stimulation of the perforant pathway. AD mice showed increased amplitude and larger spatial distribution of response after stimulation. The reason for this increased network excitability was due to impaired inhibitory neuron function, i.e. the inhibitory neurons of the molecular layer of the dentate gyrus in hippocampus were in part unable to produce action potentials, which resulted in a slower postsynaptic firing rate. Ulrich (46) added Abeta to layer V pyramidal cells of rats. In their experiments they could show a decline in inhibitory postsynaptic currents (IPSCs), attributed to postsynaptic GABA_A_ receptor endocytosis after Abeta application. In a recent study by Ren et al. (47) Abeta was found to increase excitability of pyramidal cells in the anterior cingulate cortex of mouse brain. The reason for hyperexcitability was again due to disturbed inhibitory input. Abeta seems to interact with the dopaminergic D1 receptor system. The D1 receptor regulates GABA release in fast-spiking (FS) inhibitory interneurons. By adding a D1 receptor antagonist to the cells they could reverse the effect of Abeta, increase IPSCs and decrease pyramidal excitability whereas D1 agonists had similar effects as Abeta. The underlying working model is that Abeta leads to dopamine release in dopaminergic neurons that activates D1 receptors at FS inhibitory interneurons and thus inhibits GABA release. As a consequence, the amplitude, frequency and total number of IPSPs is decreased. The instantaneous decrement of postsynaptic amplitude and frequency is also known as a toxic effect of Abeta in the glutamatergic system (48). Hence for the present modeling approach we decided to implement this Abeta dependent impaired inhibitory function. From the literature above, potential models for this disinhibiton could be either a lower IPSP amplitude or a lower firing rate or a combination thereof.

One already established drug that assesses the pathology of hyperexcitation is memantine, an N-methyl-D-aspartate (NMDA) antagonist. Memantine is recommended for the symptomatic treatment of severe AD as a mono- and combination therapy with cholinesterase inhibitors and should be also considered as possible treatment in moderate AD in the current version of the UK National Institute for Health and Care Excellence (NICE) guidelines of dementia management (49). However, normally it is considered as an alternative or addition to cholinesterase inhibitors (49). In contrast, memantine has shown in a current meta-analysis its efficacy to improve cognitive function and reduce behavioural disturbances in AD patients compared to placebo (50). The effect was particularly caused by the moderate-to-severe AD patients (50, 51) and was also observable in combination therapies with acetyl cholinesterase inhibitors, with a significant superiority for the combination of memantine and donepezil compared to any cholinesterase monotherapies (50). It therefore is also recommended as possible first-line therapy in AD (50). In our study, we will evaluate ‘virtual memantine’ interacting with the Abeta-derived hyperexcitation.

Changes in electroencephalography (EEG) are described in AD as a general and progressive slowing of brain oscillations. In AD, cognitive decline and ^18^F-fluorodeoxyglucose (FDG) PET signal decreases are linked with increased left temporal power in the delta and the theta frequency bands, whereas temporo-parieto-occipital alpha band coherence decreases and delta coherence increases (52–54). Moreover, the spatial appearance of slow rhythms and hypometabolism in FDG PET have been linked (55, 56). A recent study produced similar findings in magnetoencephalography (MEG): A global increase of theta and a frontal increase of delta were correlated with entorhinal atrophy and glucose hypometabolism (57). In summary, a global slowing has been reported for AD, in particular a shift from alpha to theta and delta activity (52–57).

As a consequence of these findings, we will focus in our modeling approach on three main aspects of AD:

1. Spatial heterogeneous Abeta distribution in the brain
2. Hyperexcitation caused by impaired inhibitory function
3. Slowing of neural frequencies

For Abeta, we propose a change in local neuronal excitability. Therefore, we construct a model of a healthy ‘standard brain’ with an averaged structural connectivity (SC) with inferred micro-scale characteristics of excitation in those areas where a deposition of Abeta is found. We will infer this information about the local distribution of Abeta from individual AV-45 (florbetapir) positron emission tomography (PET) images. AV-45 is a PET tracer which binds to Abeta (58–61). We investigate three clinical diagnostic groups of age- and gender-matched healthy controls (HC), individuals with mild cognitive impairment (MCI) and AD patients (see method section 2.1 and **Table 1**). For the simulated EEG and the underlying local neural activity frequency we expect a slowing in rhythms and particular a shift from alpha to theta activity with disease progression. Finally, we will simulate the effect of an anti-excitotoxic drug, the NMDA antagonist memantine for which we expect a reversal of the observed EEG slowing.

**Table 1.**
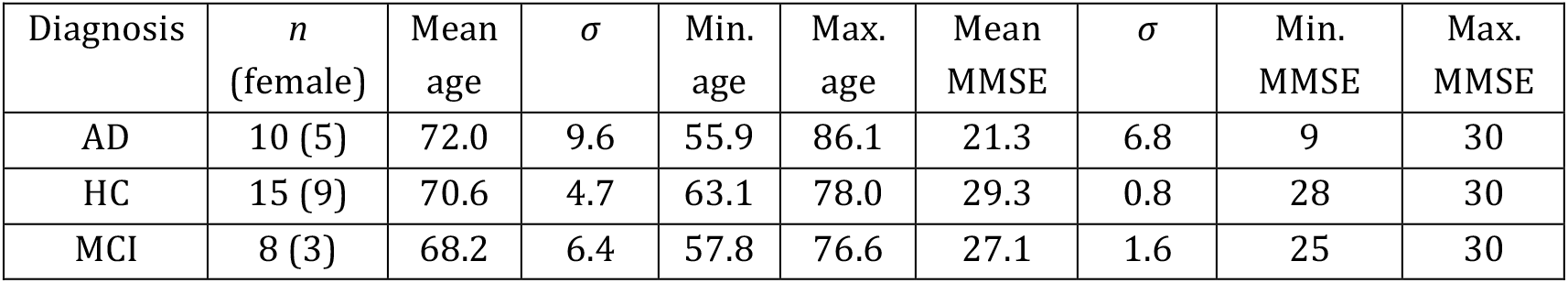
Basic epidemiological information of the study population. It is a subset of the suitable ADNI-3 participants, that had 3T imaging and all necessary image modalities. Only data from Siemens scanners was used (because this was the biggest subset of scanners).

| Diagnosis | *n*<br>(female) | Mean<br>age | $\sigma$ | Min.<br>age | Max.<br>age | Mean<br>MMSE | $\sigma$ | Min.<br>MMSE | Max.<br>MMSE |
| --- | --- | --- | --- | --- | --- | --- | --- | --- | --- |
| AD | 10 (5) | 72.0 | 9.6 | 55.9 | 86.1 | 21.3 | 6.8 | 9 | 30 |
| HC | 15 (9) | 70.6 | 4.7 | 63.1 | 78.0 | 29.3 | 0.8 | 28 | 30 |
| MCI | 8 (3) | 68.2 | 6.4 | 57.8 | 76.6 | 27.1 | 1.6 | 25 | 30 |

We will in the following provide an overview of the fundamentals of the here employed brain simulation technique. The particular strength of computational connectomics (7, 62, 63) or brain network modeling (BNM) is to unite various kinds of information in a single biophysically plausible framework (64). BNM are typically structurally informed (or constrained) by (a) geometric information of the brain, e.g. via T1 magnetic resonance imaging (MRI), and (b) the structural connectivity (SC) derived from the tractography of diffusion MRI that is supposed to represent the white matter fiber tracts (65, 66). The static three-dimensional scaffold of the brain is brought to life through the implementation of mathematical models, which generate activity at each brain region or node of the network, the so-called neural masses or population models (67–69). Population models are reduced descriptions of microscopically detailed neuronal networks (68, 70–75) - inferred for example with methods of mean field theory (76–78). They describe the so called meso-scale of the brain (79, 80), i.e. population activity as captured with imaging methods like EEG, MEG and fMRI. Some neural mass models (NMM) are linked to (and still reflect to a certain degree) neurophysiological processes at the microscopic scale while others mathematically describe the observed lumped biological behaviour not differentiating between underlying neurophysiological processes (phenomenological models). Time delays in the interaction between nodes (66, 68, 69, 81) are critical for the spatiotemporal organization of the evolving activity patterns in the brain (82, 83). Measured functional brain data such as EEG, MEG or functional MRI (fMRI) are used to tune the mathematical models – i.e. to fit selected free parameters of the model – to faithfully reproduce selected empirical features (7, 68, 76, 84–88). By performing a systematic model parameter exploration, using e.g. brute force exhaustive parameter space searches, Monte-Carlo methods or weighted optimization algorithms, we can identify the optimal parameter configuration to portray the empirical functional phenomena. Thereby, we obtain indices of the brains individual function in relation to the explored parameters. This approach opens various possibilities to not only describe dependencies (i.e., correlations), but to make statements about potential underlying causal processes, i.e. mechanisms.

In this study we used TVB, an open source neuroinformatics platform (6, 7, 68, 89) (www.thevirtualbrain.org) for large-scale BNM simulations. We have already established the software TVB, and applied it to normative datasets, stroke, epilepsy, brain tumors and neurodegenerative disease. For example, in stroke recovery, TVB models of patients were built using the patient’s structural neuroimaging data, and the dynamics of local populations were tuned to fit the patient’s functional neuroimaging data (90, 91). The obtained parameters for excitatory/inhibitory (EI) balance of local neuronal populations predicted the patient’s response to rehabilitation up to one year after therapy. Our work on epilepsy was able to infer seizure propagation with a model based on the patient’s own diffusion weighted MRI and stereotaxic EEG (92, 93). Moreover, positive surgical outcome was strongly associated with the epileptogenic zone that was excised as predicted by the patient’s TVB model. Previous work with AD patients (n = 16), controls (n = 73) and persons with amnestic MCI (n = 35), all from the Sydney Memory and Aging Study, confirms the benefit of using the model parameters to characterize cognitive status (94).

TVB provides several types of NMMs. In the present study, we selected a NMM that can simulate EEG and enables us to implement disinhibition. The wiring pattern of cortical circuitry is characterized by recurrent excitatory and inhibitory loops, and by bidirectional sparse excitatory connections at the large-scale (95). Several NMMs therefore feature projection neurons aka pyramidal cells with long axons projecting to distant cortical regions and local excitatory and inhibitory feedbacks (74, 96, 97). The NMM by Jansen-Rit comprises an elementary circuit of three interconnected NMMs (**Figure 2**) describing a cortical area (or column). It has been used to explain both epilepsy-like brain activity (98, 99) and various narrow band oscillations ranging from the delta to the gamma frequency bands (100) including intracranial EEG (69). The Jansen-Rit model has been explored extensively on a single population level (99–101) and in BNMs (85, 102, 103). The Jansen-Rit model has a rich dynamic repertoire, which was extensively described before (104).

**Fig. 2.**
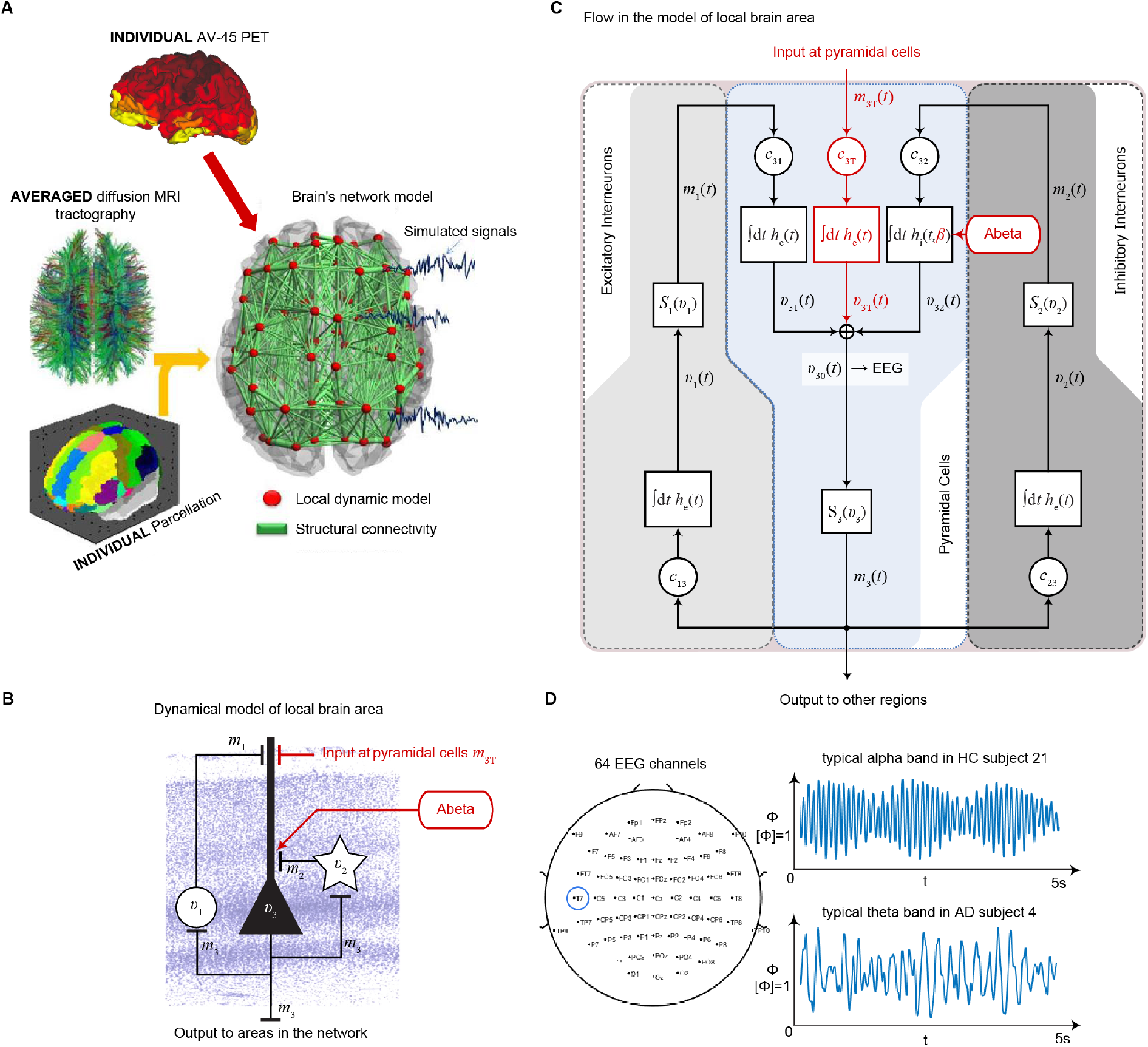
Postulated Abeta effect and its implementation to the Jansen-Rit model. **(A)** The virtual brains are based on averaged healthy connectomes and constrained by the individual regional burden of Abeta (figure modified from (62)). **(B)** In our simulation, increased excitability is caused by a disinhibition of excitatory pyramidal cells, i.e. decreased input from inhibitory interneurons. In the background a histological representation of the cortical layers: excitatory pyramidal cells (*v*_3_) and excitatory interneurons (*v*_1_) are (exemplarily) located in layer V (internal pyramidal layer), while the inhibitory stellate (inter-neurons (*v*_2_) are located in layer IV (internal granular layer). In layer I (molecular layer) we see the dendrites of the pyramidal cells, where the input from the interneurons happens. The effect to the other neuron populations is represented by m_1-3_ (background is a modified version of figure 13 from (133), license: https://creativecommons.org/licenses/by-nc/4.0/). **(C)** schematic illustration of the three interacting neural masses in the Jansen-Rit population model. The reduced inhibition is mediated by negative influence of the local Abeta burden on the inhibitory time constant *τ*_i_ (see main text for more detailed explanation). This is intended to lead to an increased activity and higher output of the pyramidal cell population. The excitatory impulse response function (IRF) is specified as *h*_e_(*t*) = *tH*_e_ exp(−t/ *τ*_e_) / *τ*_e_. The inhibitory IRF is specified as *h*_i_(*t, β*) = *tH*_e_ exp(−t/ *τ*_i_(*β*)) / *τ*_i_(*β*). These IRFs can be translated into second-order ordinary differential equations, see Eqs. 1 to 3. For explanation of the used variables, see table 2 (figure modified from (104)). **(D)** Virtual EEG as the simulation output (projection of oscillating membrane potentials to the scalp surface) reveals a shift from alpha to theta activity in AD participants. Shown is a 5 second period of exemplary EEG channel at location T7 in participant 21 (HC, above) and 4 (AD, below). The ordinate is showing the dimensionless correlate for electric potential Φ. The exemplary timeseries shows a typical simulation result in the study: in the alpha mode, which was the starting point of the Jansen-Rit model without the effect of Abeta, it produces monomorphic alpha activity with amplitude modulations (above). Mainly exclusively in the AD virtual brains a much more irregular theta rhythm appears (below).

Specifically we chose the Jansen-Rit model for the present study due to the following considerations:

1. The Jansen-Rit model comprises three interacting neural masses (representing different cellular populations) in each local circuitry: pyramidal cells, inhibitory and excitatory interneurons (**Figure 2B**). This is unique and opens the possibility to simultaneously model disinhibition, i.e. an impairment of the inhibitory neuronal subpopulation in one neural mass, and an anti-NMDAergic effect, i.e. a downscaled transmission from excitatory interneurons to pyramidal cells, at the same time.
2. The ratio of excitatory and inhibitory time constants *τ_e_* / *τ_i_* in the Jansen-Rit model is suitable to model the effect of Abeta on the inhibitory interneurons (by affecting the transmission from inhibitory interneurons to pyramidal cells, **Figure 2B-C**) and is also known to have an effect on the simulated neural frequency (104, 105).
3. Jansen-Rit can simulate physiological rhythms observable in local field potentials, (intracranially) stereo-EEG (sEEG), scalp EEG and MEG (68, 74, 104).

Our hypothesized effect of local Abeta deposition as inferred from subject-specific AV-45 PET is a decrease of local inhibition (44, 45, 47, 48, 106–108), which leads to a relatively stronger local excitation. This theory allows us translation of the Abeta distribution into the altered dynamics of a population model (**equation 6** and **Figure 3**). The hypothesized microscale (synaptic), spatially distributed effect is assumed to develop an effect at the population (mesoscale) level and to eventually propagate to the large-scale of the whole brain. A schematic illustration of this concept is provided in **Figures 1 and 2**.

**Fig. 3.**
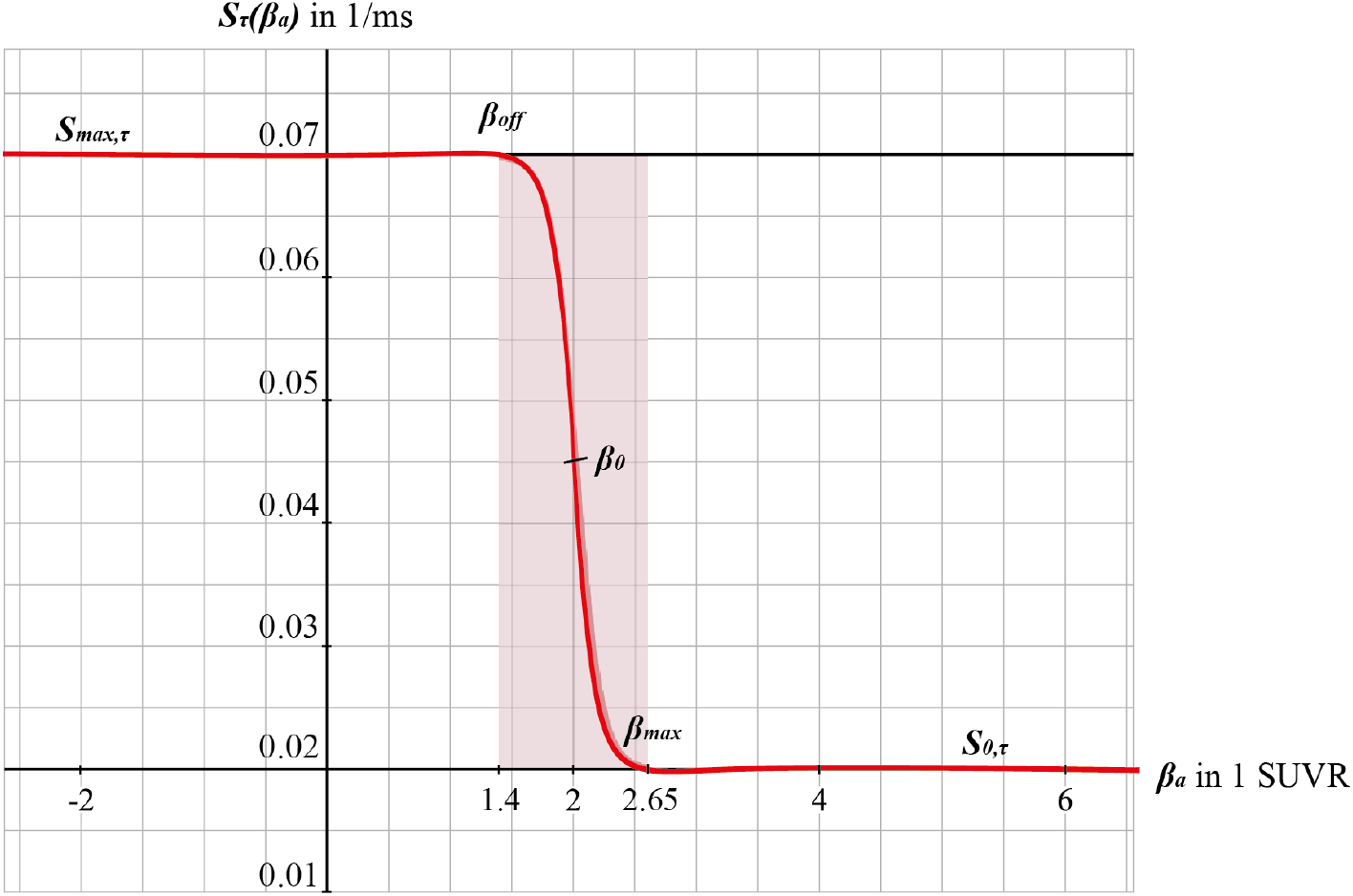
Graphs of the sigmoid transfer function of Abeta. The abscissa represents the Abeta burden *β_a_*, the ordinate represents the reciprocal *S_τ_*(*β_a_*) of the inhibitory time constant *τ*_i_. See **equation 6**.

## 2. Methods

### 2.1. Alzheimer’s disease Neuroimaging Initiative (ADNI) database

Empirical data were obtained from the Alzheimer’s Disease Neuroimaging Initiative (ADNI) database (adni.loni.usc.edu). The ADNI was launched in 2003 as a public-private partnership, led by Principal Investigator Michael W. Weiner. The primary goal of ADNI has been to test whether serial MRI, PET, other biological markers, and clinical and neuropsychological assessment can be combined to measure the progression of MCI and early AD. For up-to-date information, see www.adni-info.org.

In the presently ongoing trial, ADNI-3, the measurements contain T1, T2, DTI, fMRI, Tau PET, Abeta PET and FDG PET for the participants. The total population of ADNI-3 will contain data of about 2000 participants (comprising AD, MCI and HC, see http://adni.loni.usc.edu/adni-3/). As inclusion criterion for AD patients the diagnosis criteria of NINCDS-ADRDA from 1984 were used, which contains only clinical features (109). Inclusion criteria for both HC and MCI were a Mini Mental State Examination (MMSE) score between 24 and 30 as well as age between 55 and 90 years. For MCI in addition, the participant must have a subjective memory complaint and abnormal results in another neuropsychological memory test. To fulfil the criteria for AD, the MMSE score had to be below 24 and the NINCDS-ADRDA criteria for probable AD had to be fulfilled (109). Imaging and biomarkers were not used for the diagnosis. For the full inclusion criteria of ADNI-3 see the study protocol (page 11f in http://adni.loni.usc.edu/wp-content/themes/freshnews-dev-v2/documents/clinical/ADNI3_Protocol.pdf). An overview of the epidemiological characteristics of the participants included in this study can be found in **Table 1**.

### 2.2. Data acquisition and processing

All images used in this study were taken from ADNI-3. To reach comparable datasets, we used only data from Siemens scanners with a magnetic field strength of 3T (models: TrioTim, Prisma, Skyra, Verio). However, some acquisition parameters differed slightly. See supplementary material with **tables S1-S6** for the metadata. The following imaging modalities were included: **T1 MPRAGE**. TE = 2.95 - 2.98 ms, TR = 2.3s, matrix and voxel size differ slightly. **FLAIR**. TE differs slightly, TR = 4.8s, matrix size = 160 · 256 · 256, voxel size differs slightly. **DWI** (only for 15 HC participants to create an average healthy SC). TE = 56 −71 ms, TR = 3.4 - 7.2s, matrix size = 116 · 116 · 80, voxel size = 2 · 2 · 2, bvals = [0, 1000] or [0, 500, 1000, 2000], bvecs = 49 or 115. **Siemens Fieldmaps and PET Data** (AV-45 for Abeta). The preprocessing of imaging data can be subdivided in that of structural images, DWI and PET.

#### Structural MRI

We calculated an individual brain parcellation for each included participant of ADNI-3. We followed the minimal preprocessing pipeline (110) of the Human Connectome Project (HCP) for our structural data using Freesurfer (111) (https://surfer.nmr.mgh.harvard.edu/fswiki/FreeSurferMethodsCitation), FSL (112–114) and connectome workbench (https://www.humanconnectome.org/software/connectome-workbench). Therefore, we used T1 MPRAGE, FLAIR and filedmaps for the anatomical parcellation and DWI for tractography. This consists of a Prefreesurfer, Freesurfer and Postfreesurfer part. We skipped the step of gradient non-linearity correction, since images provided by ADNI already are corrected for this artefact. Also, the MNI templates were used at 1mm resolution instead of 0.7mm. In the Freesurfer pipeline we skipped the step of downsampling our data from 0.7mm^3^ to 1mm^3^, and all recon-all and intermediate steps were performed with the original image resolution. We then registered the subject cortical surfaces (32 000 vertices) to the cortical parcellation of Glasser et al. (115) using the multimodal surface matching (MSM, see (116)) tool. For the registration we used cortical thickness, MyelinMaps, cortical curvature and sulc from the subject and template surface. We mapped the parcellation on the surface back into the grey matter volume with connectome workbench. This volume parcellation surfed as the mask for the connectome and PET intensity extraction.

#### PET images

We used the preprocessed version of AV-45 PET. These images had following preprocessing already performed by ADNI: Images acquired 30 - 50 min post tracer injections: four 5-minute frames (i.e. 30 - 35min, 35 - 40min,…). These frames are co-registered to the first and then averaged. The averaged image was linearly aligned such that the anterior-posterior axis of the subject is parallel to the AC-PC line. This standard image has a resolution of 1.5 mm cubic voxels and matrix size of 160 · 160 · 96. Voxel intensities were normalized so that the average voxel intensity was 1. Finally, the images were smoothed using a scanner-specific filter function. The filter functions were determined in the certification process of ADNI from a PET phantom. We used the resulting image and applied the following steps: Rigid aligning the PET image to participants T1 image (after being processed in the HCP structural pipeline). The linear registration was done with FLIRT (FSL). The PET image was than masked with the subject specific brainmask derived from the structural preprocessing pipeline (HCP). To obtain the local burden of Abeta, we calculated the relative intensity to the cerebellum as a common method in the interpretation of AV-45-PET, because it is known that the cerebellum does not show relevant AV-45 PET signals and can therefore act as a reference region for inter-individual comparability between patients (58, 117). The intensity of gamma radiation, which is caused by a neutralization reaction between local electrons and the emitted positrons of the nuclear tracer is measured for each voxel in the PET image and divided to the cerebellar reference volume: the standardized uptake value ratio (SUVR). We therefore receive in each voxel a relative Abeta burden *β* which is aggregated according to the parcellation used for our present modelling approach (see below). Thus, we obtain a value *β_a_* for the Abeta burden in each brain region *a*. The cerebellar white matter mask was taken from the Freesurfer segmentation (HCP structural preprocessing). The image was then partial volume corrected using the Müller-Gärtner method from the PETPVC toolbox (118). For this step the gray (GM) and white matter segmentation from Freesurfer (HCP structural preprocessing) was used. Subcortical region PET loads were defined as the average SUVR in subcortical GM. Cortical GM PET intensities were mapped onto the individual cortical surfaces using connectome workbench tool with the pial and white matter surfaces as ribbon constraints. Using the multimodal parcellation from Glasser et al. (115) we derived average regional PET loads.

#### DWI

We calculated individual tractography only for included HC participants of ADNI-3 to average them to a standard brain template (see 2.3 below). Preprocessing of the diffusion weighted images was mainly done with the programs and scripts provided by the MRtrix3 software package (http://www.mrtrix.org).

The following steps were performed:

*Dwidenoise*. Denoising the DWI data using the method described in Veraart et al. (119).
*Dwipreproc*. Motion and eddy current correction using the *dwipreproc* wrapper script for FSL (https://mrtrix.readthedocs.io/en/latest/dwi_preprocessing/dwipreproc.html)
*Dwibiascorrect*. B1 field inhomogeneity correction using ANTS N4 algorithm
*Diw2mask*. brainmask estimation from the DWI images.
*Dwiintensitynorm*. DWI intensity normalization for the group of participants.
*Dwi2response*. The normalized DWI image was used to generate a WM response function. We used the algorithm described by Tournier et al. (120) in this step.
*Average_response*. An average response function was created from all participants.
*Dwi2fod*. Using the spherical deconvolution method described by Tournier et al. (121) we estimated the fibre orientation distribution using the subject normalized DWI image and the average response function. From the DWI data a mean-b0 image was extracted and linear registered to the T1 image. The inverse of the transform was used to bring the T1 brain masked and *aparc+aseg* image (from HCP structural preprocessing) into DWI space. The transformed *aparc+aseg* image was used to generate a five tissue type image.
*Tckgen*. Anatomical constrained tractography (122) was performed using the iFOD2 algorithm (123). Tracks in the resulting image were weighted using SIFT2 algorithm (124). We mapped the registered parcellation from Glasser back into the volume. The cortical and subcortical regions than were used to merge the tracks into a connectome.

#### EEG Forward solution in TVB

After structural preprocessing with the HCP pipeline we used the individual cortical surfaces and T1 images to compute the person specific Boundary Element Models in Brainstorm (125). Scalp, outer and inner skull were modelled with 1922 vertices per layer. Using the default ‘BrainProducts EasyCap 65’ EEG cap as locations for the signal space and the cortical surface vertices as source space. The leadfield matrix was estimated using the adjoint method in OpenMEEG with the default conductivities 1, 0.0125 and 1 for scalp, skull and brain respectively. Because we are performing region-based simulations only (i.e. no vertex-wise modelling), the leadfield matrix was simplified by summing the coefficients of vertices that belong to the same region. EEG signal was generated by matrix multiplication of the neural time series with the lead field matrix.

### 2.3. Virtual human standard brain template out of averaged healthy brains

We use the SCs of all ADNI-3 participants of the group HC, derived from the diffusion-weighted and structural MRI, to average them to one connectome matrix. Two of the HC participants included in the average template were excluded for simulations because it was impossible to compute their leadfield matrices for EEG calculation. Therefore, we use an arithmetic mean 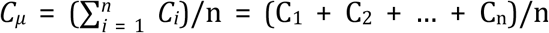, wherein *C_μ_* is the averaged SC matrix, *n* is the number of HC participants and *C_i_* ist the individual SC matrix. The SC matrix and the organization of the corresponding graph can be found in **Figure 4**. As it can be seen in **Figure 4B**, general characteristics of physiological SCs as symmetry, laterality, homology and subcortical hubs are maintained in the averaged connectome. By choosing an averaged SC instead of individual SCs, it was possible to control all factors except of the individual Abeta distribution supporting our intention to compare the simulated activity that resulted from a ‘pathogenic’ modification by Abeta.

**Fig. 4.**
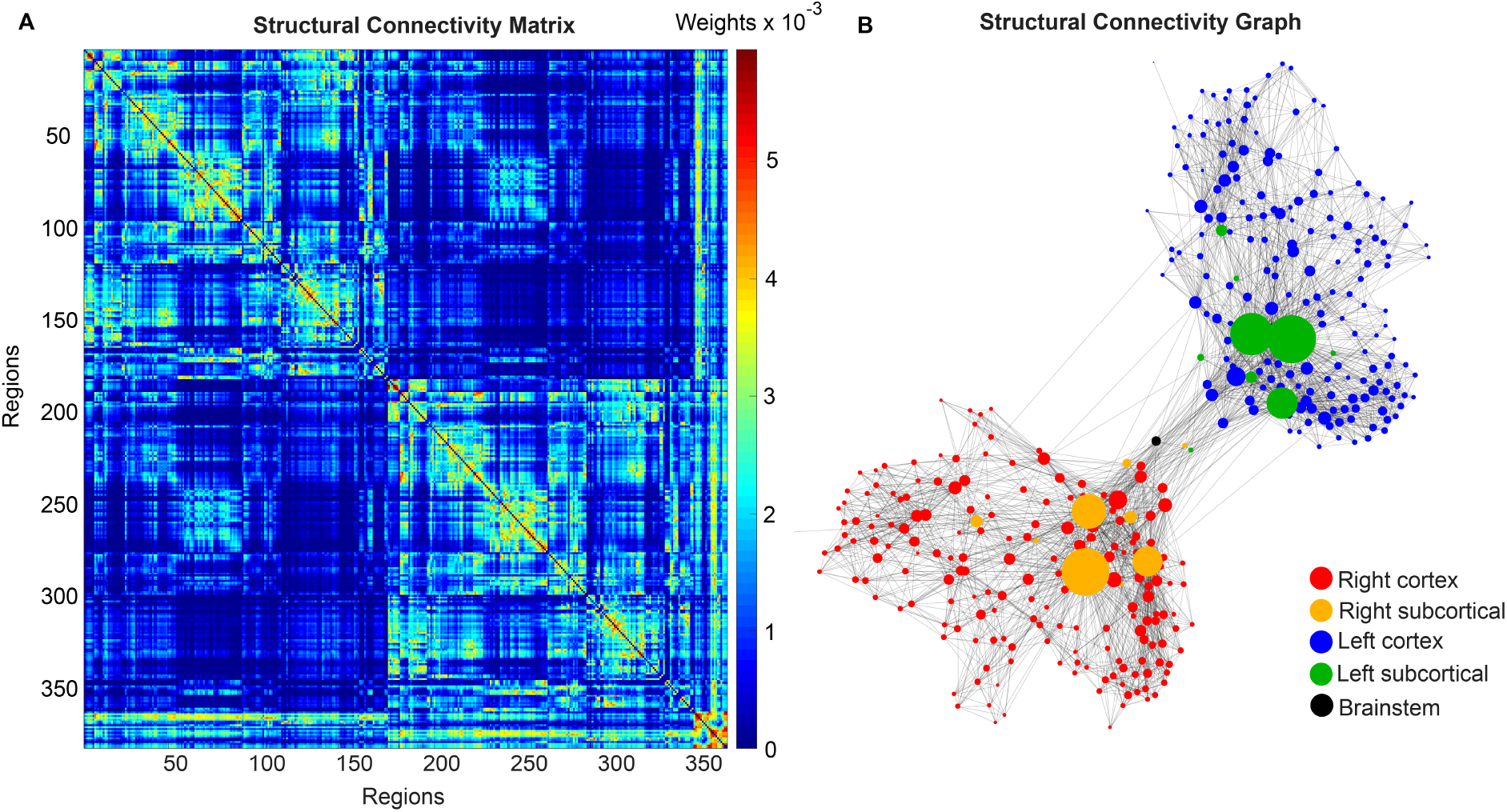
Underlying average HC structural connectome. **(A)** SC Matrix of the underlying averaged SC, showing the DWI-derived connections weights. 379 regions are in following order: 180 left cortical regions, 180 right cortical regions of Glasser parcellation, (115), 9 left subcortical regions, 9 right subcortical regions, 1 brainstem region. It gets obvious the difference between interhemispherical commissural fibers (lower weights, with a slightly pronounced diagonal between homologous regions) and intrahemispherical association fibers (higher weights). Moreover we can observe the strong connection pattern of the subcortical areas (above region 360). **(B)** Graph of the underlying SC. As a threshold, only the strongest 5% of connections were kept for binary transformation to the adjacency matrix for the graph. Node positions are derived from the inner structure of the graph by a ‘force’ method (134), assuming stronger forces and therefore smaller distances between tightly connected nodes. It can be seen that the laterality is kept in the graph structure (also for subcortical regions) and the whole graph is highly symmetric. Node size linearly represents the graph theoretical measure of structural degree for each node. Most important hubs are subcortical regions. The shown features of symmetry, laterality, homology and subcortical hubs indicate that the averaged SC still kept its physiological characteristics.

### 2.4. Cause-and-effect model of Abeta in the Jansen-Rit model

The dynamics of the Jansen-Rit model show a rich parameter dependent behavior (104). A bifurcation analysis of the single population Jansen-Rit model (in contrast to network embedded interacting populations) catalogues and summarizes the repertoire of the model. Bifurcation here refers to a qualitative change in the system behavior with respect to parameter changes. Qualitative changes can be for instance the shift from waxing and waning alpha rhythm as observed in resting human brains to spike wave discharges as observed during epileptic seizures. Bifurcation diagrams explore the qualitatively different states (divided by bifurcations, see **Supplementary Figure 1**, from (104)). The bifurcation analysis revealed an important feature of the Jansen-Rit model, which is bistability, that is, the coexistence of two stable states for a certain parameter range (i.e., regime). The bistable regime allows the coexistence of two self-sustained oscillatory states for the standard parameter configuration (74) and **Table 2** of which one state generates rhythmic activity in the alpha band and the other one produces slower big spike-wave complexes in theta rhythm. Changes in the kinetics of excitatory and inhibitory PSPs (i.e., changes of time constants) change the model behavior in a way which makes it suitable to scale, that is, to speed up or to slow down dynamics (104). The results of the systematic parameter exploration of the excitatory and inhibitory time constants is summarized in **Supplementary Figure 2**. For our study, to achieve this dynamic behavior of two limit cycles, we used first a very low input on the pyramidal cells (firing rate 0.1085/ms) and no input on the inhibitory interneurons to not overlay the Abeta effects we introduce here. Here the system operates near the subcritical Andronov-Hopf and the saddle-saddle bifurcations (leftmost region in **Supplementary Figure 1**). For the time constants, we used the area of alpha rhythm (blue area in **Supplementary Figure 2**) as control condition without any effect of Abeta. The detailed parameter settings can be found in **Table 2**.

**Table 2.** Used parameters for each Jansen-Rit element in the large-scale brain network (74).

| *Variable* | Description | Value | Unit |
| --- | --- | --- | --- |
| $H_e$ | Maximum amplitude of EPSP. Also called average synaptic gain. | 3.25 | 1 mV |
| $H_i$ | Maximum amplitude of IPSP. Also called average synaptic gain. | 22.0 | 1 mV |
| $\tau_e$ | Excitatory dendritic time constant | 10 | 1 ms |
| $\tau_i(\beta_a)$ | Inhibitory dendritic time constant as a function of Abeta load | [14.29, 50] | 1 ms |
| $v_0$ | Is the mean PSP threshold for 50% of maximum firing rate | 6 | 1 mV |
| $e_0$ | The firing rate for $v = v_0$ The maximum firing rate is $2e_0$. | 2.5 | 1 mV |
| $r_v$ | Steepness of the sigmoid transfer function. | 0.56 | 1/mV |
| $c_{31}$ | Average number of synaptic contacts from inhibitory to pyramidal cells | 108 | 1 |
| $c_{13}$ | Average number of synaptic contacts from pyramidal to inhibitory cells | 135 | 1 |
| $c_{32}$ | Average number of synaptic contacts from excitatory to pyramidal cells | 33.75 | 1 |
| $c_{23}$ | Average number of synaptic contacts from pyramidal to excitatory cells | 33.75 | 1 |
| $m^0_{3T}$ | Input firing rate at the pyramidal cells. | 0.1085 | 1/ms |
| $G$ | Global structural connectivity scaling factor | [0, 600] | 1 |
| $S_{\max, \tau}$ | Maximum value of the inhibitory rate (reciprocal of inhibitory time constant) | 0.07 | 1/ms |
| $S_{0, \tau}$ | Minimum value of the inhibitory rate (reciprocal of inhibitory time constant) | 0.02 | 1/ms |
| $\beta_{\max}$ | 95<sup>th</sup> percentile value for the Abeta burden $A\beta$ as the PET SUVR for all regions and all participants | 2.65 | 1 |
| $\beta_{\text{off}}$ | Cut-off-value for the Abeta burden $A\beta$ as the PET SUVR, from which one a pathological meaning is suspected | 1.4 | 1 |

The information about the local Abeta burden is derived from the individual AV-45 PET. As there exists no established clinical standard for SUVR cut-off thresholds differentiating normal form pathological Abeta loads. To scale the possible neurotoxic effect in a realistic way, we need to approximate at what point Abeta toxicity occurs. Following the literature, a 96% correlation to autopsy after Abeta PET was achieved via visual assessment of PET images. The corresponding SUVR cut-off was 1.2 (58). Another study showed a higher cut-off point at SUVR ≥ 1.4 for a 90% sensitivity of clinically diagnosed AD patients with an abnormal Abeta PET scan (126). We use here the higher cut-off threshold of SUVR ≥1.4. Consequently, we propose a cause-and-effect model for Abeta that is mapping molecular changes to computational brain network models:

The inhibitory time constant *τ*_i_ in each point is a function of *β_a_*. The higher Abeta SUVR, the higher is the synaptic delay and therefore *τ*_i_. We decided for this implementation via a synaptic delay because of several reasons:

1. We are focusing on disease linked alterations of EEG frequencies. Hence, we intended to assess a model feature that is already known to be frequency-effective, i.e. it can vary resulting simulated EEG frequencies. From former explorations of the Jansen-Rit-model we know that the neural frequencies are influenced by the ratio of excitatory and inhibitory time constants (104).
2. Cellular studies are supporting the hypothesis of altered inhibition, such as decreased IPSP frequency in AD (45, 47, 48) – hence we decide for mapping Abeta on the inhibitory time constant enabling for IPSP frequency modulation.
3. By using a time-effective feature, we intended to differentiate the micro-scale neurotoxic effect of Abeta on synaptic level (46–48) from connectivity-effective phenomena on a larger scale, which could e.g. be modeled by an alteration of connection strength.

We develop a transform function to implement the PET SUVR in parameters of the brain network model. Specifically, we postulate a sigmoidal decrease function that modifies the default value for inhibitory time constant *τ*_i_ (**equation 6** and **Figure 3**). We assume the healthy brain without super-threshold Abeta burden operates in a region of the parameter space, which is close to a network criticality. A criticality describes an area in the parameter space, where subtle changes of one variable can have a critical impact on others (127) (in this case bifurcations, see **Supplementary Figure 1**. The thresholding ‘cut-off’ value *β*_off_ – differentiating normal form pathological Abeta burden – was chosen according to the literature, stating that only after a certain level of tracer uptake a region is considered pathological (*β*_off_ = 1.4, see above). The maximum possible Abeta burden value *β*_max_ was chosen to be the 95% percentile of the Abeta regional SUVR distribution across all participants. The midpoint of the sigmoid was chosen such that it was half the way between *β*_off_ and *β*_max_. The steepness was chosen such that the function converges to a linear function between *β*_off_ and *β*_max_.

### 2.5. Brain Network Model construction and simulation

For the reasons stated in the above introduction, for our simulation approach we selected the Jansen-Rit model (68, 72, 74, 85, 98, 100, 101, 104, 128). The differential equations are presented in **Equations 1 - 6** (74). The employed parameter values can be found in **Table 2**.

Excitatory projections onto pyramidal cells at location *a* in discretized space (*a* = 1, 2,…, *N: N* = 379 regions):

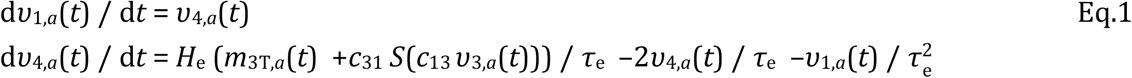

Inhibitory projections onto pyramidal cells at location *u* in space:

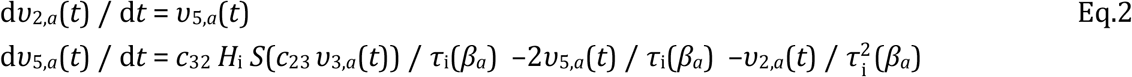

Projections of pyramidal cells *u* in space:

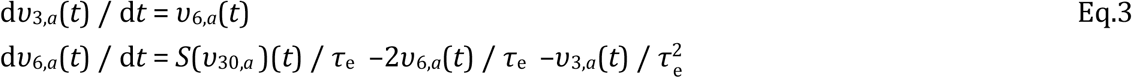

with the mean postsynaptic potential of the pyramidal cells

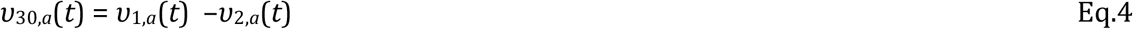

and the nonlinear potential-to-firing-rate transfer function

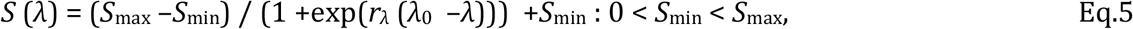

with, *λ* = *ν, S*_*ν*,max_ = 2*e*_0_ and *S*_*ν*,min_ = 0. Incoming mean firing rates *m*_3T,*a*_(*t*) at the pyramidal cells at location *a* from other brain regions *b* = 1, 2,…, *N*, where *N* is the number of 379 regions are given by 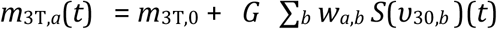, where *m*_3T,0_ is baseline input *m*_3T,0_ = const. for ∀*t* and all locations ∀*a*. The global coupling factor *G* scales the connections *w_a,b_* incoming at location *a* from all *b* provided by the SC. In all populations, the state variable [*ν*_1_, *ν*_2_, *ν*_3_]_*a*_(*t*) are the mean membrane potentials and the derivatives thereof with respect to time *t*, namely [*ν*_4_, *ν*_5_, *ν*_6_]_*a*_(*t*) represent the mean currents.

To model how the local Abeta load *β_a_*, measured by the Abeta PET SUVR is affecting the inhibitory time constant we introduce the following transfer function (**Figure 3**):

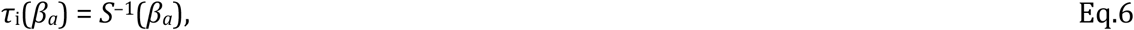

with *r_β_a__* = 2ln(*S*_max_ · 1000 ms – 1) / (*β*_*a*,off_ – *β*_*a*,max_) and *β*_0_ = (*β*_*a*,off_ + *β*_*a*,max_) /2. Note that the inverse of the time constant *τ*_i_ is the rate. The Abeta load affects the inhibitory rate following a sigmoid curve. The rate ranges between *S*_min_ and *S*_max_ and the time constant ranges consequently between 1/*S*_max_ and 1/*S*_min_.

To simulate the model using TVB, physical space and time are discretized. The system of difference equations is then solved using deterministic Heun’s method with a time step of 5 ms. We used a deterministic method to avoid stochastical influences since the simulation was performed in the absence of noise.

The system was integrated for 2 minutes and the last 1 minute was analyzed in order to obtain the systems’ steady state and diminish transient components in the time series due to the initialization.

We explore a range of 0 ≤ *G* ≤ 600 which provides an overview about the possible population level behaviors at different states of network coupling. Because the coupling factor *G* has a crucial influence on the external input on the neuronal populations, this allows different regions to operate in different dynamical regimes, as it can be seen in the bifurcation diagrams of **Supplementary Figure 1**. Global coupling factor *G* that was sampled between *G* = 0 (i.e., isolated regions) and *G* = 600 with a step size of Δ*G* = 3. The initial values were taken from 4000 random time points for each state variable in each region. The length of 2 minutes for the simulations was chosen with the aim to diminish possible transient components due to the initialization of state variables at *t* = 0. For analysis we used only the second minute of the simulated signals. No time delays are implemented in the large-scale network interactions since they are not required for the emergence of the here evaluated features and setting them to zero increases reduces required computation resources.

### 2.6. Spectral properties of the simulated EEG

In TVB, we simulate EEG as a projection of the oscillating membrane potentials inside the brain via its electromagnetic fields to the skin surface of the head (68) using the individual lead field matrices which take into account the different impedances of white matter, grey matter, external liquor space, pia and dura mater, the skull and the skin. Our lead-field matrices considered the impedances of three compartment borders: brain-skull, skull-scalp and scalp-air (7, 66, 129, 130). The postsynaptic membrane potential (PSP) considered for the projection is the one of the pyramidal cells, as they contribute the mayor part to potential changes in EEG (131). The PSP is calculated by summing the synaptic input from excitatory and inhibitory subpopulations to the pyramidal cells. The baseline PSP was derived as the mean PSP across time for every region. For the LFP or EEG peak frequency, we computed the power spectrum using the ‘periodogram’ function of the Scipy python toolbox (132). From the spectrogram the ‘dominant rhythm’ was identified as the frequency with the highest power.

## 3. Results

### 3.1. Abeta-inferred dynamics lead to individual spectral patterns

We analysed the dominant frequency in the simulated EEG and regional neural signal (referred to as local field potential (LFP) (**Figure 5G – 5J**).

**Fig. 5.**
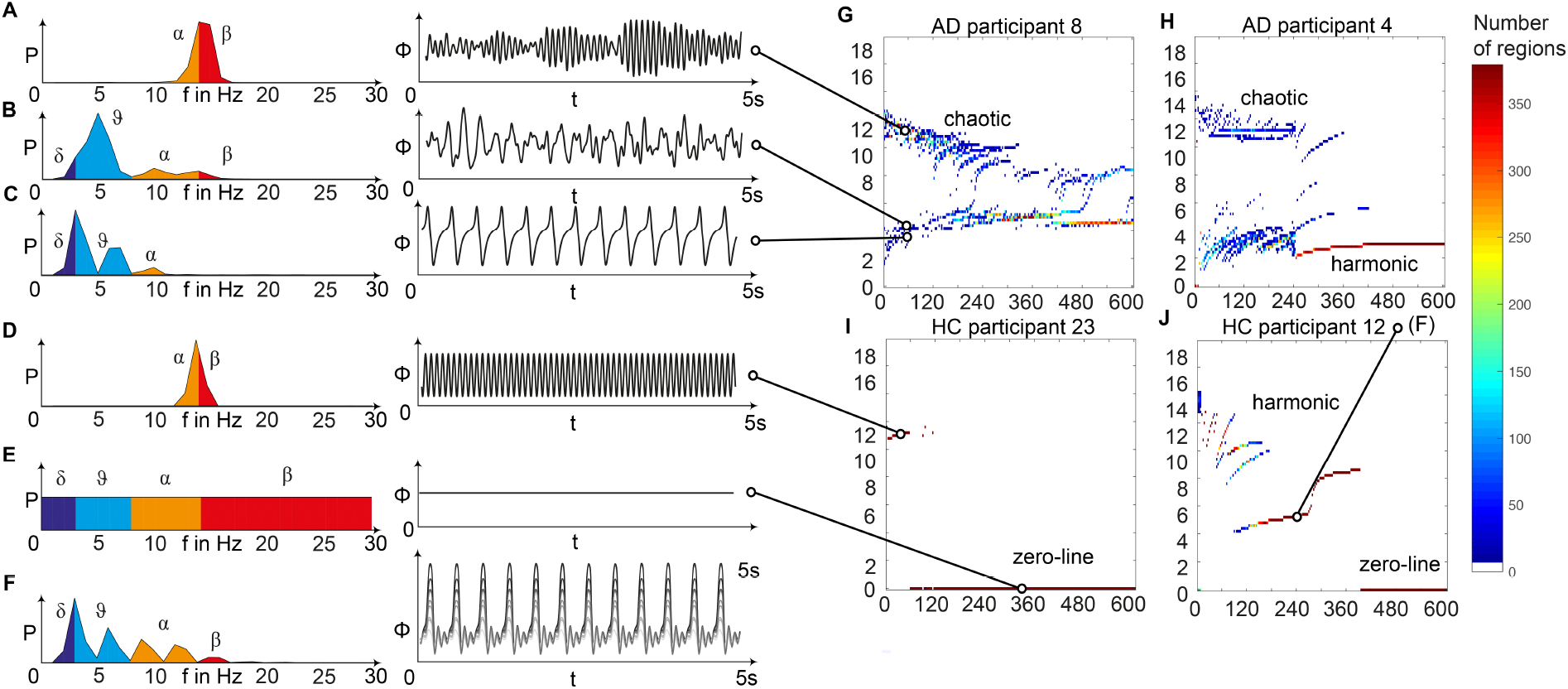
Spectral behavior in individuals of the different groups. **(A, B, C, D, E, F)** – selected timeseries and spectrograms. On the left: power spectral density of neural activity for an exemplary region with different values for G. Abscissa is frequency, ordinate is an estimate of the spectral power (dimensionless equivalent of amplitude per 1 Hz). Colors are representing the EEG frequency bands from delta to beta, indicated with greek letters (note that this is regional neural activity, not EEG). Corresponding time series on the right: neural activity at single regions, each showing 5 seconds. Abscissa is time, ordinate is a dimension-less equivalent of the electric potential. **(A)** shows an irregular, amplitude modulated alpha to beta rhythm, **(B)** an irregular theta with some delta and alpha inside. In **(C)** we can observe a monomorphic spike signal with a theta/delta frequency of 3 Hz and higher order harmonies. **(D)** Shows a monomorphic (high) alpha rhythm, **(E)** shows the zero-line with a continuous power spectrum. **(F)** Time series of 10 regions in a G area of hypersynchrony. We can see here the synchronized signals in theta rhythm and multiple harmonies of higher order in the spectrogram. **(G, H, I, J)** – four exemplary participants with different types of frequency behaviors along the range of coupling. Shown are the regional simulated dominant frequencies (y) along global coupling G (x) for individual exemplary participants 4, 8, 12 and 23. See **Supplementary Table 7** in supplementary material for participant IDs. Color indicates the density of regions with the same coordinates. The sources of the timeseries on the left **(A-F)** are marked in the plots. **(G)** irregular or chaotic rhythm with two clusters in alpha and theta. AD participant 8. **(H)** chaotic behavior for lower G, then harmonic and hypersynchronization. AD participant 4. **(I)** early zero-line, with monomorphic alpha activity at very low G. HC participant 23. **(J)** harmonic to zero-line rhythm, with a G area of hypersynchrony in alpha and theta, depending on G.

We observed a physiologically looking irregular behavior with two frequency clusters in the alpha and in the theta spectrum (**Figure 5G**). This behavior is expressed in the area of lower global coupling G for all 10 AD participants and in 3 out of 8 MCI and 4 out of 15 HC participants. The irregular time series and the broad continuous frequency spectra (**Figure 5B**) of network regime in 0 < *G* < 150 are indicative for deterministic chaos. Such chaotic network regimes in a BNM have already been reported using the same local dynamic model (Figure 2 in (85)). Beside this emerging chaotic behavior in our simulations other phenomena occurred in the parameter space exploration: a state of hypersynchronization between regions (**Figure 5H, 5J**) and a state of a ‘zero-line’ with no oscillations that clearly does not reflect a physiological brain state (**Figure 5I, 5J**).

In order to locate the individual simulations in the spectrum of possible dynamics, meaning in the range of possible Abeta load, we examined extreme values of Abeta distribution. The virtual brains with a mean Abeta load of zero (**Supplementary Figure 3A**) and with the maximum Abeta load at all regions (**Supplementary Figure 3B**), we see as expected for the Abeta-free system a behavior similar to the low-Abeta-containing HC participants. This is not surprising, because when the HC subjects do not have a high Abeta signal, the dynamics will converge to those with zero Abeta, which is in fact then only determined by the underlying standard SC and therefore remains the same for all participants. However, the homogeneous application of maximum Abeta burden does not lead to an AD-like pattern but shows a zero-line at the whole spectrum.

To give a mathematical explanation of those phenomena, we related each participants Abeta-burden to the corresponding inhibitory time constant τ_i_ and used former analyses of the uncoupled local Jansen-Rit model (104) to estimate the bifurcation diagrams for the coupled system in this study (**Figure 6**). Shown diagrams allow to predict and explain the occurrence of alpha and theta rhythms or zero-lines depending on the underlying Abeta burdens. The variation of τ_i_ by local Abeta burden fundamentally influences the systems bifurcations by shifting the bifurcation point along the range of external input to the pyramidal cells. As a consequence, different values of Abeta lead to a variable occurrence of two limit cycles and a stable focus. Therefore, for a single region with constant external input on pyramidal cells, depending on Abeta the region might be in an alpha limit cycle, in a theta limit cycle, in a bistable condition where both cycles are possible or in a stable focus.

**Fig. 6.**
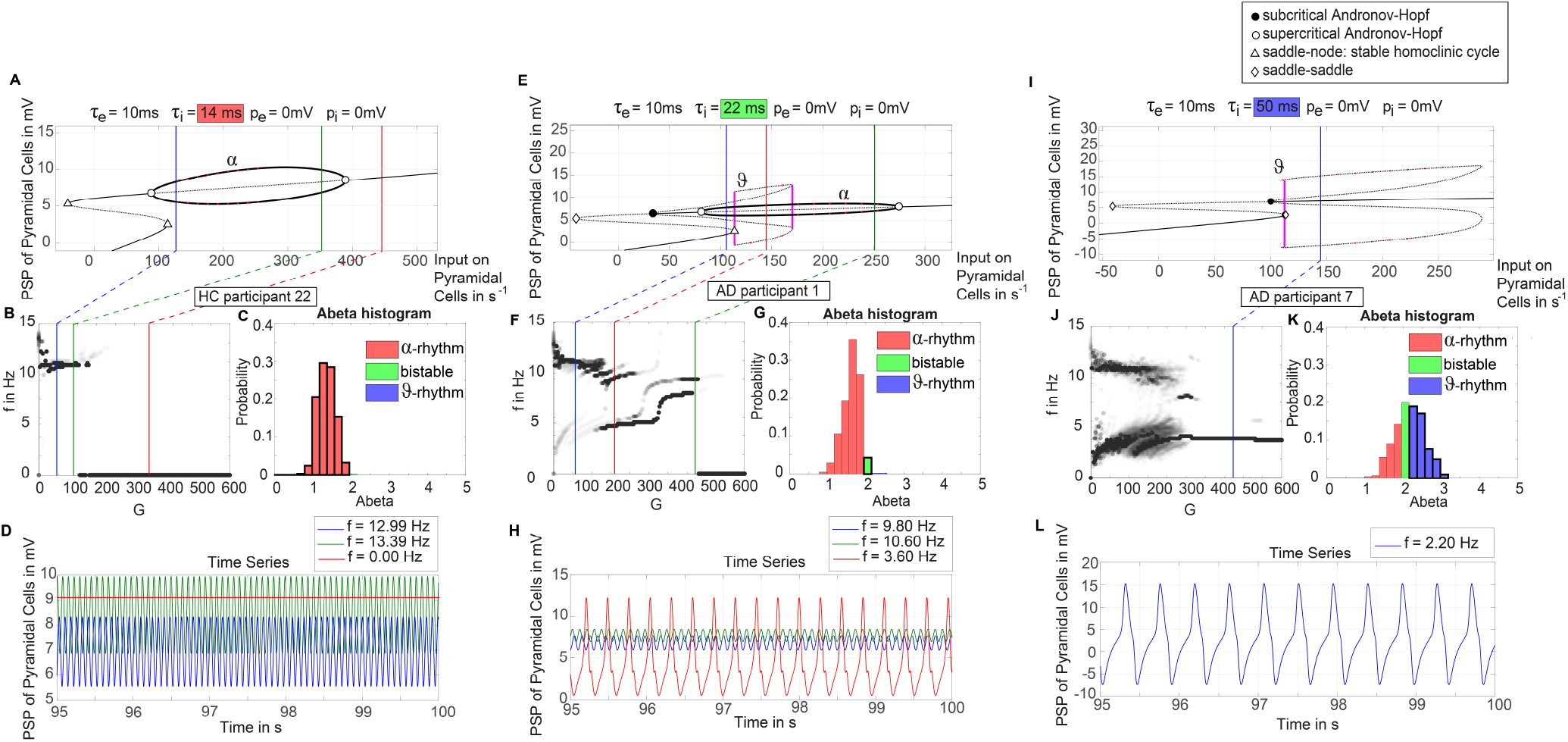
Exemplary bifurcation diagrams of the Jansen-Rit model for three different inhibitory time constants linked to three different local Abeta burdens. The modulation of the inhibitory time constant *τ*_i_ by Abeta induces shifts in the corresponding bifurcation diagrams. All bifurcation diagrams **(A, E, I)** show the postsynaptic potential of pyramidal cells (y) depending on the pyramidal input (x) for uncoupled simulations (104). The default input on pyramidal cells starts at a firing rate of 108.5/s. Because of the potential-to-firing-rate transfer function (**equation 5**), global scaling factor G is not only affecting the input currents, but also the firing rates. For higher values of G, the input on pyramidal cells is expected to increase. First Columns, panels **(A, B, C, D):** Bifurcation diagram with the default time constant of 14ms. This appears in the simulation if the Abeta SUVR is below the clinical cut-off 1.4, because then the time constant is unaffected according the transfer function in **equation 6**. In this situation, there is only one limit cycle existing, which produces a frequency in alpha range **(A)**. After increasing the input on the pyramidal cells, the alpha cycle collapses and transforms to a stable focus, where no oscillations appear in the absence of noise. This is the ‘zero-line’ in our results. **(B, D)**: HC participant 22 shows monomorphic alpha for lower G (green and blue line) and zero-line for higher G (red line). The distribution of regions with this dynamical regime is shown in (C): almost all regions of participant 22 are in this ‘alpha regime’ with an inhibitory time constant between 14ms and 20ms (red columns in **(C)**) This homogeneity explains the low variance of rhythms shown in the lower G ranges of **(B)**, because all regions are in the same limit cycle and in the absence of artificial noise there is no possibility for an amplitude modulating factor. Second column, panels **(E, F, G, H):** Bifurcation diagram with a time constant of 22ms, which corresponds to an intermediate Abeta load and a bistable dynamical regime which occurs for time constants between 20ms and 28ms. **(E)** Starting at the blue line (initial condition in alpha cycle), with an increased input on the pyramidal cells (e.g. by the network) it gets possible to reach the second limit cycle, which produces a theta rhythm and coexists with the alpha cycle while the pyramidal input is in a specific range (120/s – 170/s). When the input is increased too much (e.g. by many connections of the network or by increased coupling factor G), the theta cycle disappears and the system jumps back to the alpha cycle and later on to the stable focus, which shows no oscillations in the absence of noise. This can explain some of the spectral behaviors we observed typically in the AD group **(F, H)**: It starts with chaotic rhythms in alpha (blue line) and theta (red line) and in the shown AD participant 1 then gets synchronized to either alpha or theta. With higher couplings, the frequency gets more probably synchronized to alpha (green line), because higher G indicates a higher pyramidal input and therefore a higher attraction of the alpha cycle. **(G)** Remarkably for the shown participant is the fact that the bistable behavior is caused by a very small amount of regions in bistable regime, which propagate the theta rhythm to most other regions in the area 200 < G < 300. Third column, panels **(I, J, K, L):** Bifurcation diagram with a time constant of 50ms, which correlates to a 95^th^ percentile Abeta load and above. Those high Abeta burdens lead to a theta dynamical regime, which occurs for time constants between 28ms and 50ms. In comparison to **(E)**, the alpha limit cycle disappeared in **(I)**. Therefore, we expect only theta rhythms or an activity at the stable focus. The theta cycle now begins shortly above the initial condition of pyramidal input without the alpha cycle in between. For an initial input of 108.5/s the system is in a stable focus. This may explain why in the simulation with maximum Abeta load at all regions (so each with a time constant of 50ms) we see a zero-line without alpha at lower G values (figure S1 in the supplementary material). **(J, L)** A state of theta-only rhythm appeared in few AD participants at higher Gs (blue line). In the spectral behavior of AD participant 7, we can moreover observe a strong bistable pattern with chaotic frequency distributions for G < 300. This is likely caused by the high amount of bistable regions **(K)**, while the synchronization to theta in higher G is an effect of the high proportion of regions in theta regime.

### 3.2 Simulated EEG slowing in AD is caused by heterogeneous Abeta distribution

**Figure 7** displays how the mean dominant rhythms differ between the groups. In the range below G = 100 we find a slowing in the AD group. Since in the range of lower G all three groups exhibit realistic frequency spectra and no zero-lines we consider this range of G as ‘physiological’. Significant differences appear between AD and non-AD for ranges of high and low G and also for high alpha and low theta rhythms (**figure 7**). The heterogeneous distribution of Abeta (in contrast to an averaged homogeneous distribution) plays a crucial role in the development of this AD-specific slowing. This is indicated by simulations with the mean averaged Abeta of each participant mapped on all regions. The simulations revealed a regionally more homogenous behavior in all groups (Supplementary material, **Supplementary Figure 4**). Moreover, with homogeneous distribution of Abeta the slowing in AD participants does not appear: we don’t see a significant change in the theta band (**Figure 7B**). This is a strong indicator for the importance of the individual Abeta distribution and a proof for the necessity of heterogeneous excitotoxic effects for the creation of neural slowing.

**Fig. 7.**
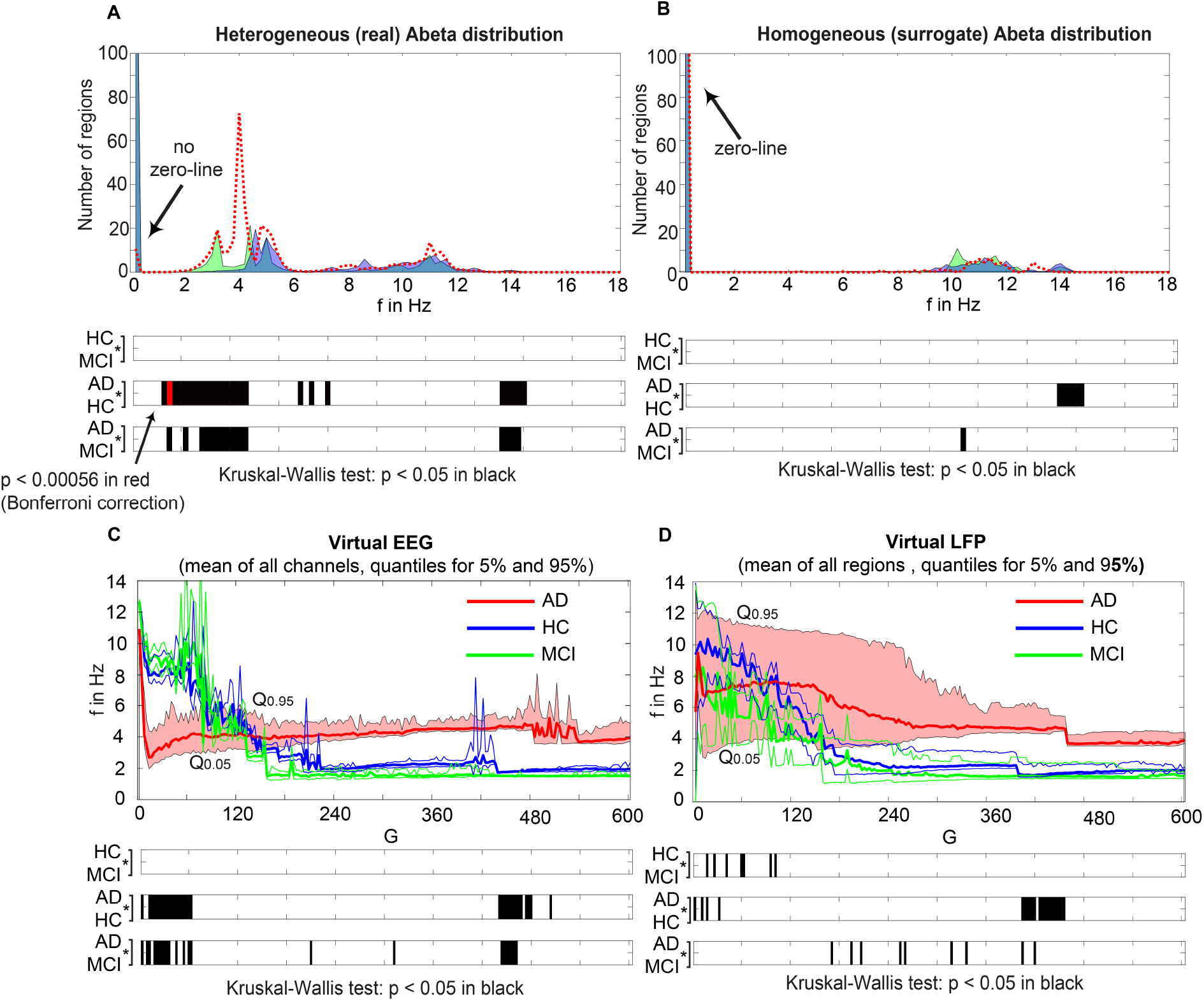
AD-specific slowing in EEG and PSP and influence of the heterogeneous pattern of Abeta distribution to the spectral behavior. **(A, B)**: The panels show the ‘spectrograms’, more precise the amount of regions with a dominating frequency averaged for all G values and the subjects of each group. Below, black bars are indicating significant differences for all 90 examined frequencies by a Kruskal-Wallis test (compared were the means of the amount of regions in each group having this particular frequency). In **(A)**, for the empirical Abeta distribution pattern, the red dotted line (AD) diverges from the non-AD participants with a strong presence of dominating theta (peak at 4 Hz) and the absence of zero-line rhythm (except of very few regions, see arrow). Significant differences only appear between AD and each HC and MCI, namely for high alpha / low beta and for theta/delta (black bars). At f = 1.2 Hz (red bar), the significance level is also achieved when using a strict Bonferroni correction (p < 0.05 / 90). In contrast, **(B)** shows the same plot if the spatial distribution was ‘blurred’: There is no visual difference between the behavior of the three groups, and also no theta rhythm is existing in the simulations. All groups have a dominating zero-line behavior averaged across the full G range (see arrow). However, there are some frequencies that significantly differ between AD and each HC and MCI in alpha / beta range, which could be also visually related to small peaks at the plots beside. In theta and delta, where we would expect to see the slowing, there is no significant difference at all. Due to readability, for **(A)** and **(B)** the y-axis was limited to the amount of 100 regions. In **(A)**, the zero-line peak of HC and MCI ends at 211, in **(B)** all zero-line peaks end at 323. The different spectra lead to different G-dependent mean frequencies for the groups, which significantly differ in areas of high and low G: **(C, D)** – comparison of EEG and LFP between groups. Mean dominant rhythms across all simulated EEG channels **(C)** and region-wise simulated LFPs **(D)** for all analyzed global coupling values. The frequencies of AD patients are significantly different in EEG as well as in the regional neuronal population signal. Filled shapes and thin lines represent the quantiles at 0.95 and 0.05 for each group. **(C)** for EEG one can see that the 95%-quantile of AD and HC as well as MCI is not overlapping in the physiological area of lower G, where AD tends to slower frequencies. In a Kruskal-Wallis test, the difference between the means of all channel frequencies per subject in the three groups is significant for AD and non-AD at 0 < G < 60 (each AD to HC and AD to MCI: p < 0.0001). They are also significantly different in the area of higher G, where AD is faster -at 450 < G < 470 (each AD to HC and AD to MCI: p < 0.0001). **(D)** for simulated regional neural signal the slowing effect is less prominent. The broader range of frequencies for AD is represented by the high and low limit of the 95%-quantile. This can be related to the two frequency clusters in AD at alpha and theta, which are not frequently apparent in non-AD (as in **Figure 5**). In a Kruskal-Wallis test, the difference between the means of all regional frequencies per subject in the three groups is only continuously significant for AD against HC at 400 < G < 450 (AD compared to HC p < 0.0001). For the other comparisons, only isolated G values deliver significant differences in the area of low G (HC and MCI) and intermediate G (AD and MCI). Because of the big amount of tests necessary to test all global coupling values, none of the tested G values achieved Bonferroni corrected significance. However, because we assume that neither the frequencies at **(A)** and **(B)** nor the G values at **(C)** and **(D)** are independent variables (which is also the reason for the ‘grouped’ clusters of significance at alpha and theta and G = 50 and G = 450), a Bonferroni correction is not necessary.

### 3.3 Intra-individual ratio of high versus low Abeta burden across all regions determines simulated EEG frequency spectrum – distinct spatial configurations of Abeta do not matter for slowing

We next examined how LFP/EEG slowing is related to the underlying Abeta burden (**Figure 8**). We revealed significant linear dependencies for all groups between Abeta burden and frequency. We found a strong inverse dependency for AD (R^2^ = 0.625), i.e. an Abeta-dependent EEG slowing. In contrast, for non-AD participants the relation was revers, i.e. higher values of Abeta caused EEG acceleration.

**Fig. 8.**
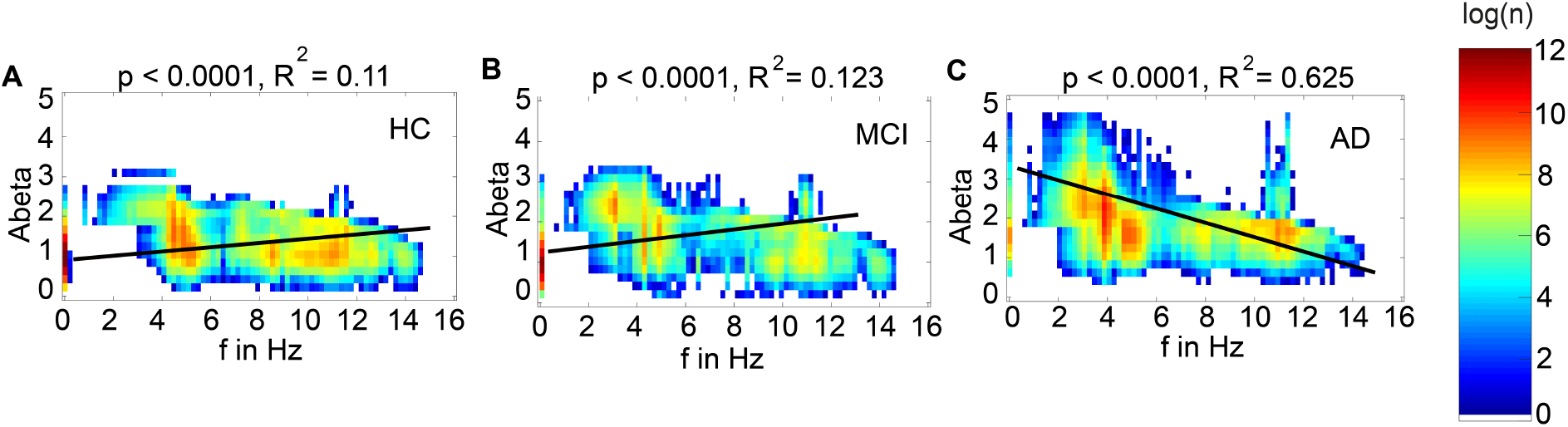
Abeta-dependent slowing of LFPs is specific for AD participants. Meanwhile there is a significant linear dependency between Abeta and LFP frequency for all groups, only for AD a higher burden of Abeta leads to a decrease of frequency. HC and MCI show inverse correlations. Plotted are density plots showing the dependency between the local Abeta loads and LFPs. **(A)** HC group, **(B)** MCI group and **(C)** AD group. The matrices are containing the resulting regional peak frequencies for all examined coupling values G for all participants. Linear regressions (black lines) revealed highly significant regression coefficients (p<0.0001). A strong linear dependency between mean Abeta and LFP, that explains the greater part of the variance, is only apparent in the AD group **(C).** 37.5% of the variance yet cannot be explained by this linear dependency. Moreover, only for AD the dependency leads to slower frequencies for higher Abeta SUVRs, meanwhile HC and MCI have slightly faster frequencies for higher Abeta SUVRs. Visually one can see at least four contributing patterns in the AD group **(C)**: (1) the linear decrement of frequency for higher Abeta, shown by the regression line, (2) the two frequency clusters (orange spots) at alpha and theta, (3) some regions with the zero-line behaviour, particular those with low Abeta (thin line at the left, with SUVR of about 1.5), and (4) a broad variability of frequencies for regions of the same Abeta SUVR (horizontal distribution). These phenomena cannot be explained completely by a linear dependency and moreover not by a linear system at all. The criticality that divides the dynamics into three different frequency modes (zero, alpha and theta) is a phenomenon of the Jansen-Rit model as a non-linear system (**Figure 6** and **Supplementary Figure 6**) and the broad frequency distribution is (probably) a network effect.

To test if specific regions are more important for the observed phenomena, we had to overcome the bias that only specific regions were strongly affected by Abeta. I.e., for the empirical Abeta distribution we cannot say e.g. for a region with high Abeta if it shows EEG/LFP slowing only because of its high Abeta value or because of its specific spatial and graph theoretical position in the network. Therefore, we next performed simulation with 10 random spatial distributions of the individual Abeta PET SUVRs for the 10 AD participants. In these simulations, the neural slowing appeared similarly to the empirical spatial distributions of Abeta (**Supplementary Figure 5**), which indicates a minor role of the distinct spatial patterns of Abeta. Instead, the ratio of regions corresponding to the three different dynamical regimes (alpha, theta and bistable) determined the simulated frequency spectrum (**Supplementary Figure 6**). For an optimal value of G with 100 < *G* < 150, the ratio of regions with an Abeta value in theta regime best corresponded to the ratio of regions with theta frequency in LFP. Moreover, the number of regions in different regimes enables to predict the individual spectral behaviour across G.

The results of random spatial distribution of Abeta PET SUVRs were also used for a parameter space exploration (**Figure 9**). The analysis reveals that (1) alpha rhythms are only apparent for low time constants with *τ*_i_ < 30ms, but for the full spectrum of G, more probable for lower G values; (2) relevant amounts of bistable rhythms are only apparent for 17ms < *τ*_i_ < 39ms and *G* > 120; (3) theta rhythms are present across almost the full spectra of G and *τ*_i_, with an equal appearance across G, but with a local minimum at *τ*_i_ ≈ 18ms, where the system is dominated by alpha and bistable rhythms. This exploration demonstrates two major insights. First, it confirms the crucial role of *τ*_i_ for the appearance of alpha or theta rhythms as we expect it out of the (non-coupled) bifurcation diagrams of **Figure 6**. Network effects are present (e.g. there are theta rhythms for low values of *τ*_i_), but play a minor role here. Second, the value of G does not significantly affect the probability of theta rhythm, except of an alpha-theta shift for low *τ*_i_ < 20ms and higher G > 160. This is caused by the coexistence of stable focus in alpha regime and theta limit cycle in theta regime for high pyramidal input (**Figure 6A and 6I**).

**Fig. 9.**
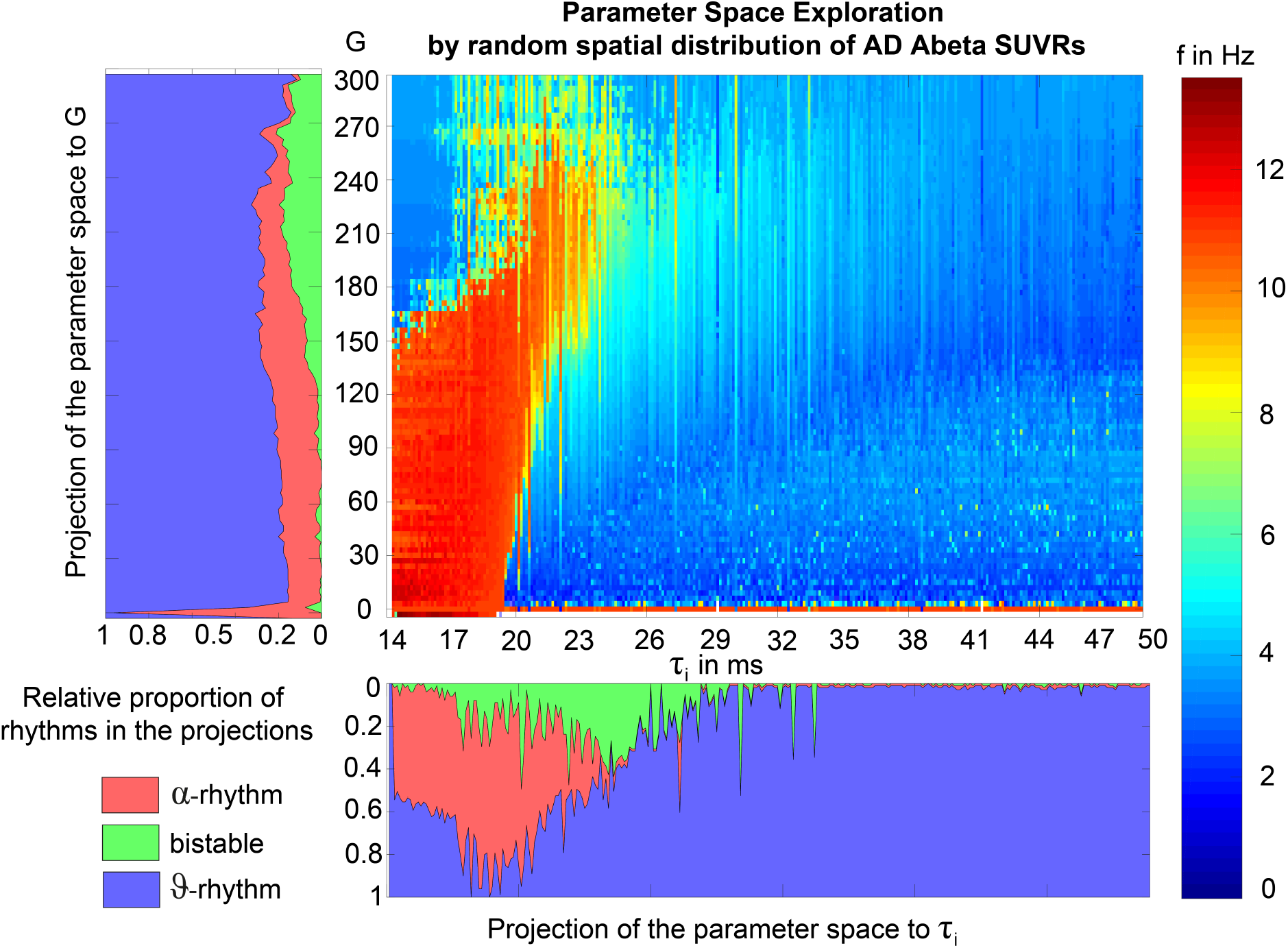
Alpha and bistable rhythms only appear in a specific part of the parameter space between G and τ_i_. This parameter space exploration was done by coupled simulations and therefore includes network effects. Frequency (by color) is presented dependent on global coupling G (x) and inhibitory time constant τ_i_ (y). Projections to G and τ_i_ are shown beside the matrix plot, here the frequencies are classified into alpha rhythm (f > 8 Hz), theta rhythm (f < 5 Hz) and bistable rhythms (5 < f < 8). No relevant proportion of zero-lines appeared in the simulations. The difference to empirical EEG classes (with slightly lower borders for theta, meaning more exactly a theta/delta rhythm) are reasonable here because of the knowledge of only two different limit cycles in the examined configuration of the Jansen-Rit model (**Figure 6**). This is also the reason for the classification of frequencies between 5 and 8 Hz as bistable. The exploration was non-systematically performed by using all regions of random distributed Abeta SUVR values of the 10 AD participants, with 10 iterations of randomization per participant. However, except single values of τ_i_, the full spectrum of τ_i_ could be explored. Single empty columns are filled with neighbor columns for better readability. In principle wee the an ‘isle’ of alpha for low coupling and low time constant, while the rest of the dynamics is dominated by theta and delta. A full frequency spectrum (also green and yellow colors) is only apparent near the borders of the alpha isle in higher coupling.

### 3.4. Neural slowing propagates to central parts of the network independently of the spatial Abeta distribution

In the analysis of spatial distribution in relation to the organization of the underlying SC network (**Figure 10**), it can be seen that unless Abeta is distributed more peripherally, the Abeta-dependent effect of neural slowing is focused to central parts of the network. Even a random distribution of Abeta SUVRs leads to this effect (**Figure 10 E-F**), indicating that this is a network effect. Probably this phenomenon is caused because the slowing effects are not only affecting the region itself, but also its local circuitry and neighbored regions. Hubs with a high degree and many close neighbours are therefore more probable of being affected by slow rhythms propagated by other regions. To relate this to empirical facts: We know from our data (**Figure 10A**) that Abeta is not deposited in hubs, but more in peripheral regions of the networks. This shows, however, how the consecutive pathologic slowing effect is afterwards focused to central and important parts of the networks. A weak peripheral affection of the inhibitory system therefore disturbs the full system seriously.

**Fig. 10.**
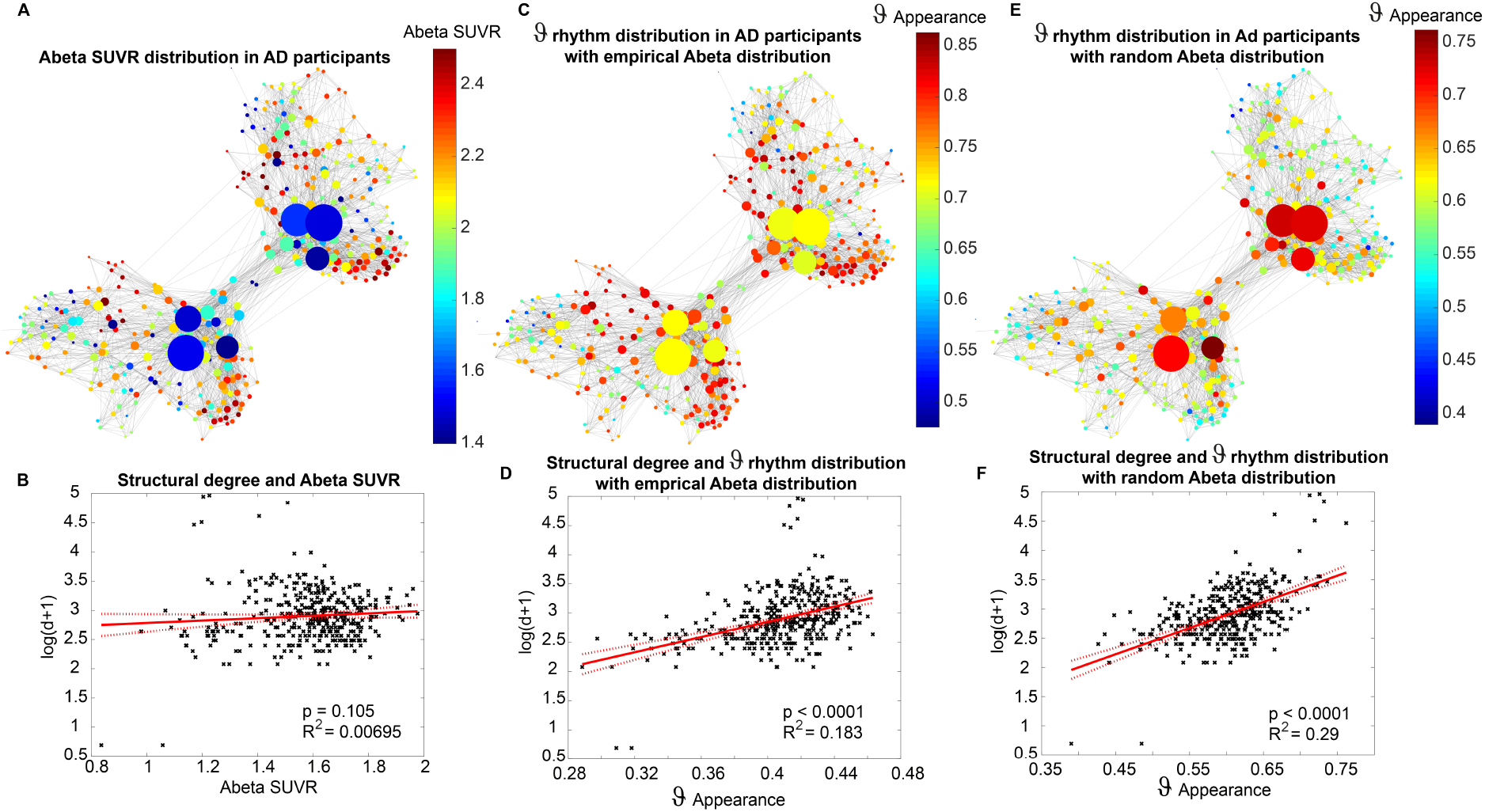
Theta rhythms affect central parts of the network independently of the spatial distribution of Abeta. **(A)** Abeta PET SUVR for AD participants: the distribution is diffuse along the cortex with no strong affection of subcortical hubs. This well corresponds to the neocortical stage C of Abeta distribution (11, 135, 136). **(B)** There is no linear dependency between the Abeta SUVR and the structural degree, as the graph above already indicates. In contrast to that, **(C)** shows the distribution of theta rhythm, computed as the proportion of each regions simulations (201 for different values of G for 10 subjects) with dominant theta rhythm (here simplified as a frequency that is below 8 Hz and not zero, so more precise the theta-delta-band). The patterns are not consistent with those of **(A)**. This indicates that not the distinct region affected by Abeta is crucial, but more its local circuitry. Moreover, one can observe that regions with a higher degree often have a high appearance of theta rhythm **(D)** and show a linear dependency with R^2^ = 0.183, in contrast to the distribution of Abeta **(B)**, which hasn’t shown such a dependency. This phenomenon is stable also for the random spatial distribution of Abeta SUVRs **(E)**: Here we see even a stronger dependency (R^2^ = 0.29) between structural degree and theta rhythm **(F)**. This is remarkable because (unless the spatial distribution is random) the ‘pathologic’ theta is focused on the hubs. This indicates that there must be network effects which concentrate the appearing theta to those regions with higher degree.

### 3.5. Virtual Therapy with the NMDA antagonist memantine

The former analyses have shown that Abeta-mediated simulated hyperexcitation can lead to realistic changes of simulated brain imaging signals in AD such as EEG slowing (**Figures 5 and 6**). We therefore wanted to know if an established way to protect the brain of the hyperexcitation, which is the NMDA antagonist memantine, can lead to functional reversibility.

The idea in our model is now that in theory memantine acts anti-excitotoxic via its NMDA antagonism and should therefore be able to weaken the hyperexcitation we introduced to the system by Abeta **(Figure 11)**. As mentioned above, the local coupling parameter *c*_32_ represents the main part of the glutamatergic transmission and can therefore also be seen as a surrogate of NMDAergic transmission. We homogeneously increased the default value of *c*_32_ stepwise to observe the effects on the system. In **Figure 11A** and **11B** one can see that it would be not useful to decrease *c*_32_ to a lower level then 0.6, because then the system does not have enough energy to produce network activity in the area of low coupling. The weakened intrinsic NMDAergic coupling has to be anticipated by a stronger global coupling. This can also be seen when the global coupling reaches high values (**Figure 11C**): the red curves of AD patients with and without virtual memantine are converging. The virtual memantine leads to a partial reversibility of the altered dominant frequencies in AD compared to HC/MCI. Virtual memantine increases the mean dominant EEG frequency. These simulated functional effects do not imply reversibility of neurodegeneration, but they illustrate how pharmacological intervention can theoretically counteract those processes. This observation provides first a potential mechanistic explanation of the pharmacodynamics of memantine. Second, it shows that TVB in general and the Abeta-hyperexcitation model of this study in particular are able to test the efficacy of treatment strategies such as drugs and have therefore the potential to be used for the discovery of new treatment options. Finally, it supports the concept of this study, where the impaired inhibitory function is modelled by an increased synaptic delay. The effects of altered delay of GABA transmission can be reversed by adjusting NMDA transmission at another subset of the local population model. This illustrates that theoretically an alteration of the inhibitory transmission dynamics may lead to disinhibition causing hyperexcitation in downstream populations, which is reversible by reduction of excitatory input.

**Fig. 11.**
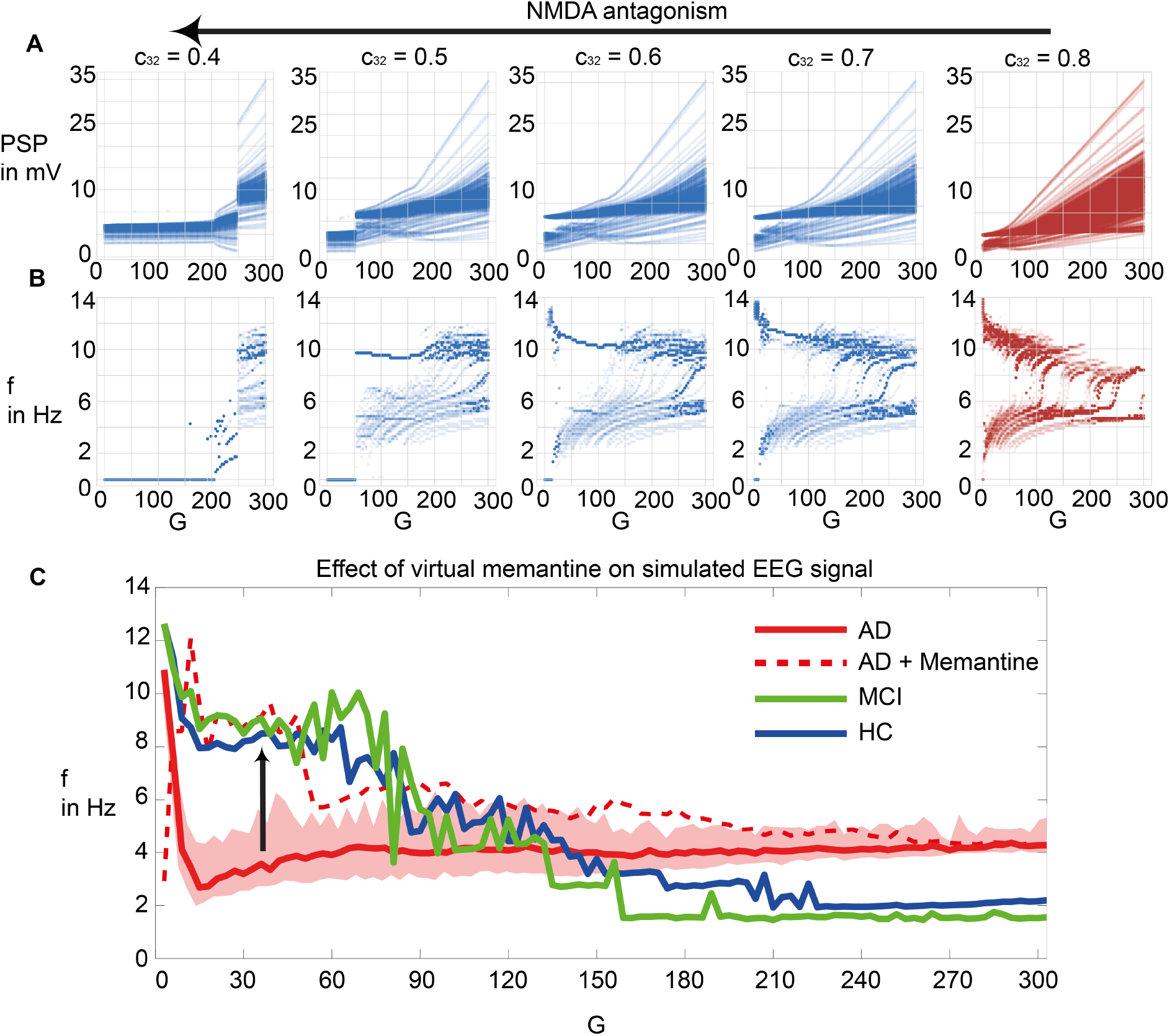
Modeling NMDA antagonism by virtual memantine. We modified the local dynamics for the AD group by homogeneously decreasing *c*_32_, which represents the coupling from excitatory population to the pyramidal cell and therefore is a surrogate for NMDA receptor activity. **(A)** PSP and **(B)** distribution of regional frequencies along different global couplings for various values for *c*_32_. The plots are similar to those in **Figures 5 and 7**. On the right, in red, we see the default value of *c*_32_ = 0.8 as in the simulations before. When scaling the factor down, one can see in **(A)** that the system needs higher global coupling to increase energy in the PSPs. When decreasing *c*_32_ too much, this leads to a dying out of activity in the area of lower global coupling, which begins at a value of *c*_32_ = 0.5. We therefore used for memantine the reduction by 0.25% when changing *c*_32_ from 0.8 to 0.6. **(C)** Mean EEG frequency for the three groups HC (blue), MCI (green), AD (red, with shadowed area for the range between 5^th^ and 95^th^ percentile) and AD with memantine (red dotted line). The virtual application of memantine shifts the AD group to the level of HC and MCI (arrow) and out of the variance of AD without memantine.

## 4. Discussion

Local Abeta-mediated disinhibition and hyperexcitation are considered candidate mechanisms of AD pathogenesis. In TVB simulations, the molecular candidate mechanism has led to macro-scale slowing in EEG and neural signal with a particular shift form alpha to theta previously observed in AD patients (52–57). These observations cannot be directly inferred by the hyperexcitation implemented in our model. Because we standardized all other factors and used a common SC for all simulations, this approach enables to examine the effects of disinhibition on an individual whole-brain level but without any other confounding factors.

1. We showed that the slowing in simulated EEG and LFP is specific for the AD group (**Figures 7 and 8**). This offers an explanation, how the shift from alpha to theta, that is observable in EEG of AD patients (52–57), could be explained on a synaptic level – namely by an impaired inhibition. This computational modelling result supports the findings of specific toxicity of Abeta to inhibitory neurons (46–48).
2. We demonstrate the computational principles underlying this Abeta dependent slowing of EEG/LFP (**Figure 6 and Supplementary Figure 6**). Dependent on the Abeta burden alpha, theta or bistable regime develop caused by an alteration of the inhibitory time constant that leads to changes of the systems bifurcation behavior (**Figure 6, Supplementary Figures 1, 2 and 6**).
3. The simulated LFP/EEG slowing in AD patients crucially depends on the spatially heterogenous Abeta distribution as measured by PET – the slowing disappears when using a homogenously distributed mean Abeta burden instead for simulation (**Figure 7**). To exhibit the slowing effect few regions with high Abeta burden are required – while the specific location of these regions seems not to be relevant (**Supplementary Figure 5**).
4. Independently of the location of high Abeta burdens in the simulated brain, slowing emerges at the core, i.e. hubs of the structural connectome (**Figure 10**). This indicates that that central parts of the system are impacted functionally by the Abeta burden. Moreover, it shows that while Abeta is often distributed in peripheral parts of the structural connectome, its functional consequences affect the important hubs. This could provide a possible explanation why a peripheral distribution of Abeta leads to severe disturbances of cognitive function.
5. We also showed that the drug memantine that is known for improving brain function in severe AD can be modeled by a decreased transmission between the excitatory interneurons and the pyramidal cells and is able to achieve a ‘normalized’ brain function *in silico*, too (**Figure 11**). This moreover demonstrates the potential of TVB to test and develop new treatment strategies.

In this study, we present proof of concept for linking molecular changes as detected by PET to large-scale brain modeling using the simulation framework TVB. This study therefore can work as a blueprint for future approaches in computational brain modeling bridging scales of neural function. For the research on AD pathogenesis, this study provides a possible mechanistic explanation that links Abeta-related synaptic disinhibtion at the micro-scale to AD-specific EEG slowing. In general, our study can be seen as proof of concept that TVB enables research on disease mechanisms at a multiscale level and has potential to lead to improved diagnostics and to the discovery of new treatments.

## Acknowledgements

Data collection and sharing for this project was funded by the Alzheimer’s Disease Neuroimaging Initiative (ADNI) (National Institutes of Health Grant U01 AG024904) and DOD ADNI (Department of Defense award number W81XWH-12-2-0012). ADNI is funded by the National Institute on Aging, the National Institute of Biomedical Imaging and Bioengineering, and through generous contributions from the following: AbbVie, Alzheimer’s Association; Alzheimer’s Drug Discovery Foundation; Araclon Biotech; BioClinica, Inc.; Biogen; Bristol-Myers Squibb Company; CereSpir, Inc.; Cogstate; Eisai Inc.; Elan Pharmaceuticals, Inc.; Eli Lilly and Company; EuroImmun; F. Hoffmann-La Roche Ltd and its affiliated company Genentech, Inc.; Fujirebio; GE Healthcare; IXICO Ltd.; Janssen Alzheimer Immunotherapy Research & Development, LLC.; Johnson & Johnson Pharmaceutical Research & Development LLC.; Lumosity; Lundbeck; Merck & Co., Inc.; Meso Scale Diagnostics, LLC.; NeuroRx Research; Neurotrack Technologies; Novartis Pharmaceuticals Corporation; Pfizer Inc.; Piramal Imaging; Servier; Takeda Pharmaceutical Company; and Transition Therapeutics. The Canadian Institutes of Health Research is providing funds to support ADNI clinical sites in Canada. Private sector contributions are facilitated by the Foundation for the National Institutes of Health (www.fnih.org). The grantee organization is the Northern California Institute for Research and Education, and the study is coordinated by the Alzheimer’s Therapeutic Research Institute at the University of Southern California. ADNI data are disseminated by the Laboratory for Neuro Imaging at the University of Southern California.

Petra Ritter acknowledges the following funding sources: H2020 Research and Innovation Action grants 826421 and 650003 and 720270 & 785907 and ERC 683049; German Research Foundation CRC 1315 & 936 and RI 2073/6-1; Berlin Institute of Health & Foundation Charité, Johanna Quandt Excellence Initiative.

Further we acknowledge Lea Doppelbauer and Jan Roediger for their helpful discussions.

## Author Contribution Statement

All authors have made substantial intellectual contributions to this work and approved it for publication. LS and PR had the idea to this study. LS, PT, ANDS and PR developed the concept and study design. LS wrote the manuscript, conducted the analysis and interpretation of results and developed the figures. PT performed the MRI and PET image processing and supercomputer simulations. PT, ANDS, MD, ANAS, VJ, ARM and PR contributed to the interpretation of the results, figure development and writing of the manuscript.

