## Supplementary Materials for "Linking molecular pathways and large-scale computational modeling to assess candidate disease mechanisms and pharmacodynamics in Alzheimer’s disease"

**SUPPLEMENTARY MATERIAL.**
**FIGURES.**

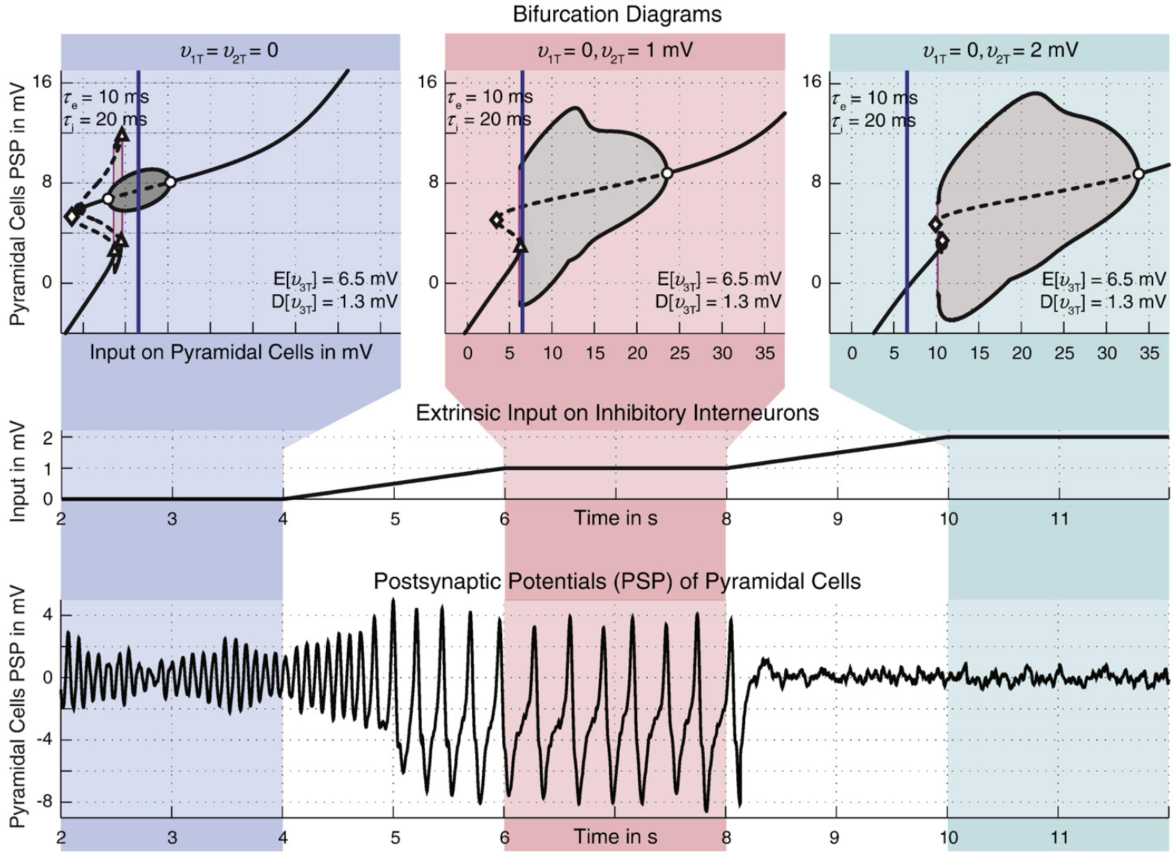


**Supplementary Figure 1. Input on pyramidal cells in Jansen-Rit model leads to fundamental changes (criticalities) in the behavior of the system.** Taken from Spiegler et al., figure 13 of (1) with permission. Input on inhibitory cells is changed with time (middle row). Shown are three different states of the model: amplitude-modulated alpha activity (highlighted in blue), slower spiking activity (red) and noise in the absence of intrinsic oscillations (green). In the top row, the corresponding bifurcation diagrams are shown for different input values on inhibitory cells. The diagrams show the bifurcations between input on pyramidal cells (x) and the PSP of pyramidal cells (y). Criticalities lead to fundamental changes of the temporal behavior of PSP for only slightly different inputs. In this case, the criticalities of the non-linear system are bifurcations in its mathematical meaning (2). Different types of bifurcations can be seen: sub-critical (black) and supercritical (white) Andronov–Hopf bifurcations, saddle–saddle bifurcations (diamonds) and saddle-node bifurcations (triangles), global bifurcations (red lines). For a more detailed description and for the context of this figure in the corresponding study, please consider the original publication (1).

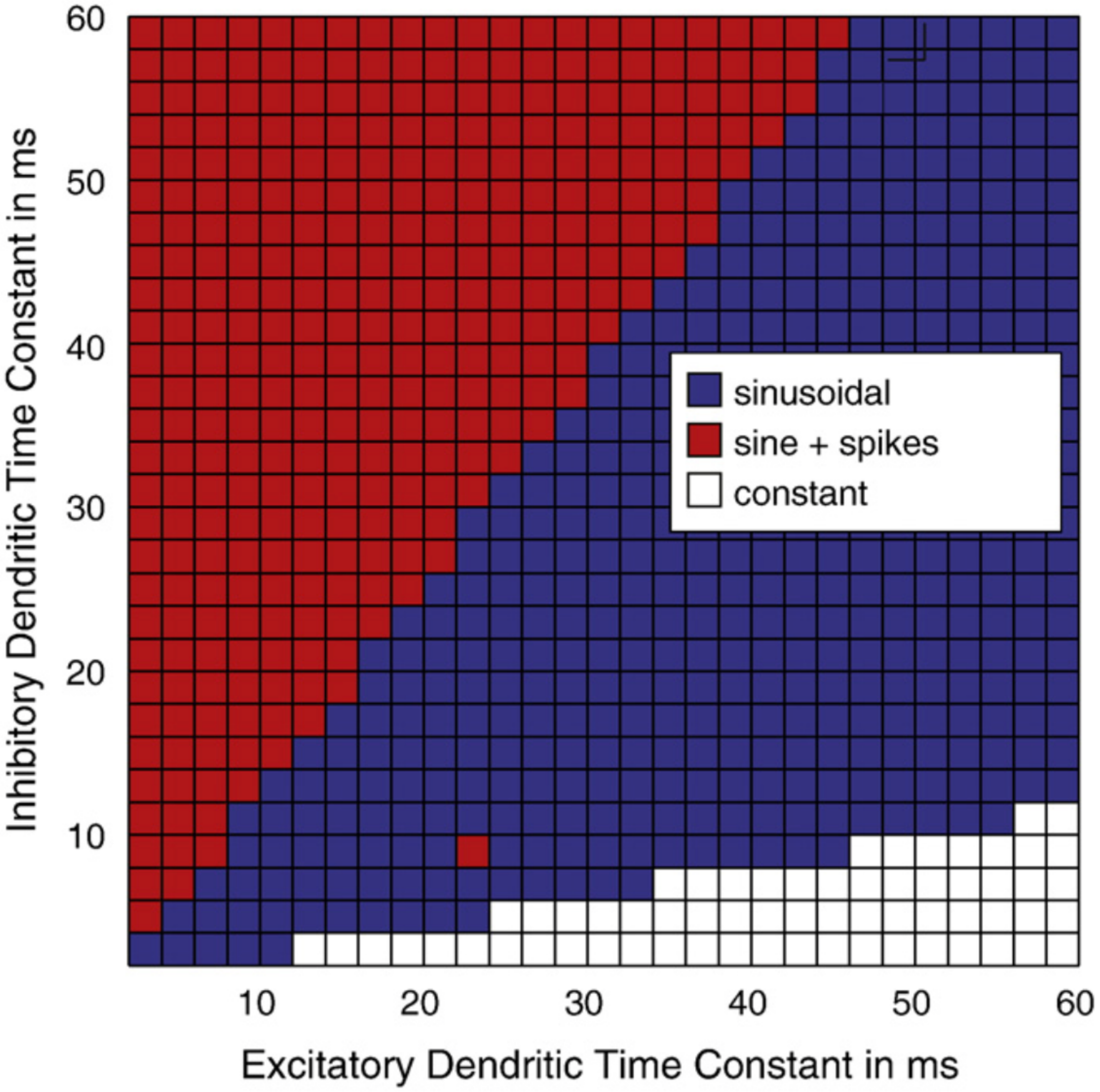


**Supplementary Figure 2. Ratio of excitatory and inhibitory time constants modulates frequency in Jansen-Rit model** Taken from Spiegler et al., figure 7 of (1) with permission. Blue regions in the parameter space show fast sinusoidal oscillations in alpha rhythm. In red areas, also slower spikes in theta rhythm get possible. For a more detailed description and for the context of this figure in the corresponding study, please consider the original publication (1).

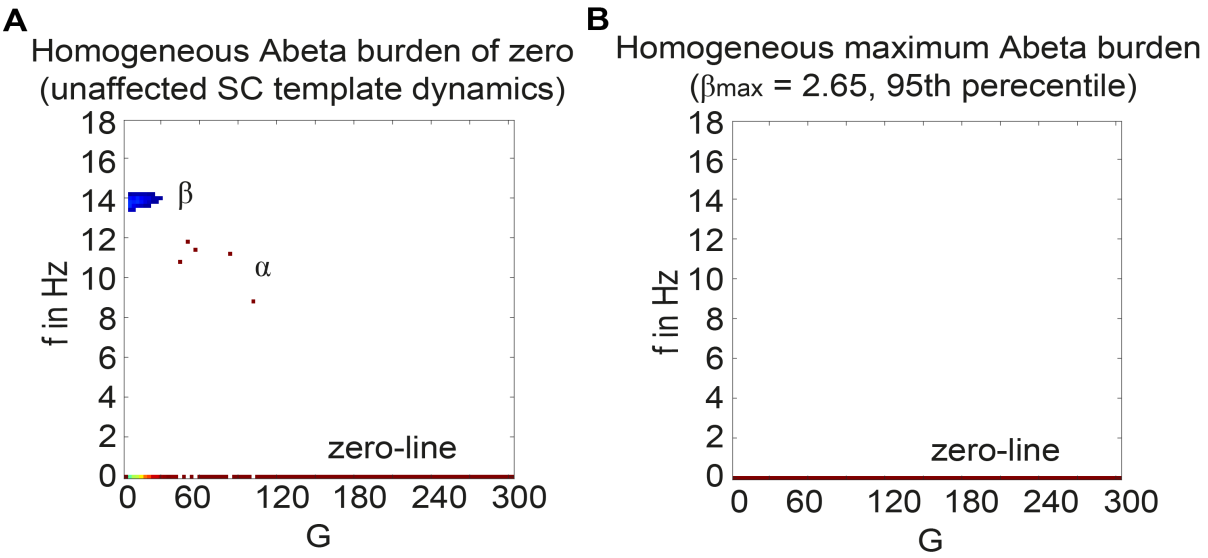


**Supplementary Figure 3. Control simulations for the standardized SC template with extreme values of Abeta burden.** Because the only individual feature in this study is the Abeta distribution, all participants show the same behaviour for an equal and homogeneous Abeta distribution. Density of regions with a specific dominating frequency at each G. Shown are the sums of all participants regions. **(A)** Homogeneous distribution with Abeta burden of zero at each region, therefore representing the unaffected dynamics that are only driven by the underlying averaged healthy SC template. There is a small beta cluster and single G values between 50 and 100 with alpha rhythms, but the behaviour is dominated by the zero-line. See also **Figure 6A**. **(B)** Homogeneous distribution with Abeta burden of $\beta_{max}$ = 2.65 at each region. Maximum Abeta was calculated by the 95$^{th}$ percentile of all regions in all participants. It is also represented in the sigmoid curve in **Figure 3**. The full G spectrum is characterized by the zero-line. See also **Figure 6I**.

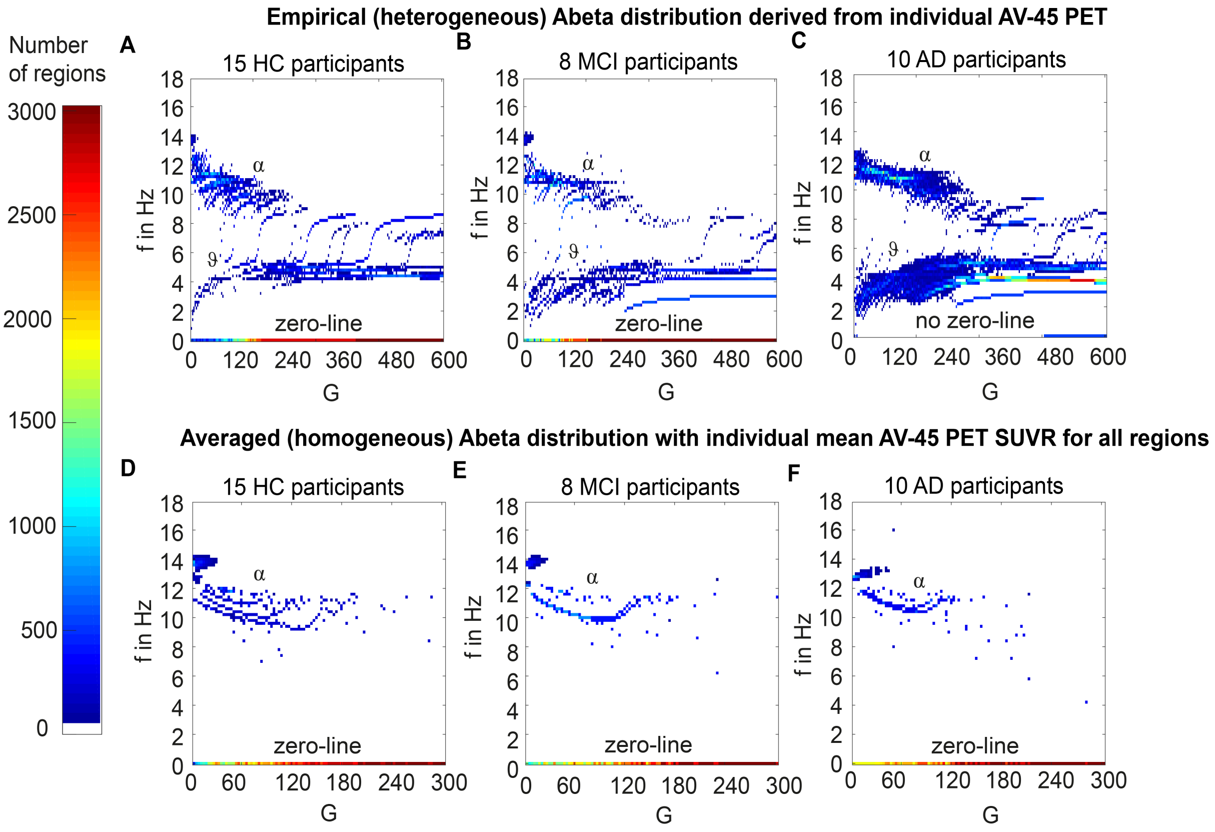

**Supplementary Figure 4. AD-specific slowing in neural frequencies and influence of the heterogeneous spatial pattern of Abeta distribution to the spectral behavior. (A, B, C) -** Overview of the different frequency behaviors dependent on global coupling factor G, summarized for the three groups HC, MCI, AD**.** The plots show the density of regions with a specific dominating frequency at each G. Shown are the sums of all participants` regions for **(A)** HC (n=15), **(B)** MCI (n=8) and **(C)** AD (n=10). Notably some regions reside at a frequency of 0 beginning at low G values for HC and MCI. This behavior is not apparent for AD, there is only a blue line (meaning low density) at zero at a high G **(C)** – because only 1 out of the 10 AD participants showed a zero-line behavior. **(D, E, F)**: corresponding plots for simulations with homogeneous distribution of averaged Abeta load for each subject, so without spatial information about Abeta distribution and without Abeta heterogeneity. The mean AV-45 PET SUVR for each participant was applied to every region of the brain. The phenomena found in panels **(A-C)** do not appear in **(D)** HC, **(E)** MCI and **(F)** AD. Namely all show a similar synchronized alpha rhythm at low G and convert early to the zero-line behaviour. This strongly supports the hypothesis that not the general burden of Abeta is the driving factor for the observed phenomena in this study. It seems to be more the spatial heterogeneity in the brain, meaning that in the brain coexist areas with different local dynamics that are influencing each other.

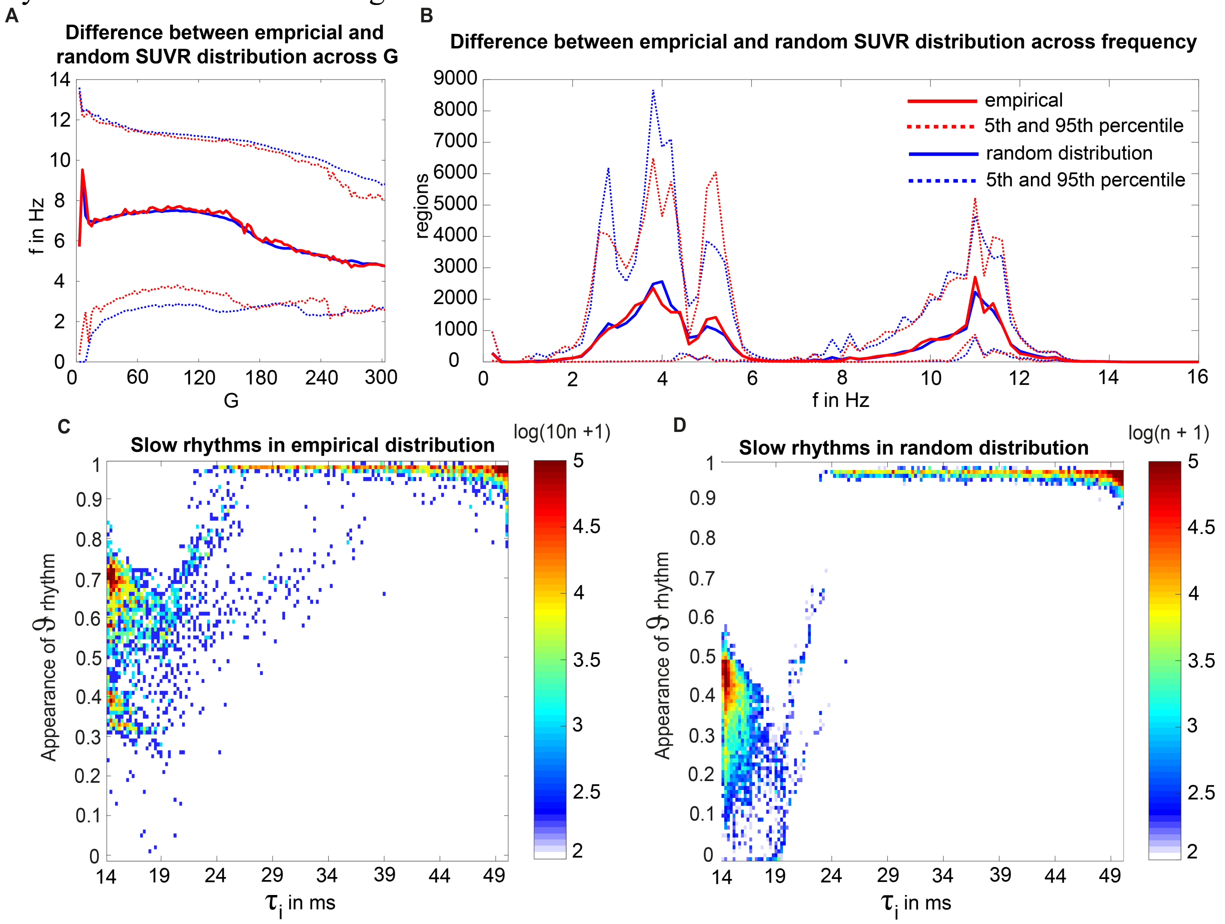


**Supplementary Figure 5. Results for random spatial distribution of Abeta PET SUVRs of AD participants. (A)** Mean dominating frequencies (y) along global coupling (x), averaged across regions and subjects. Color legend shown in **(B)**. There is no relevant difference except the smother contour of the blue line because of 10-times more simulations. **(B)** Spectrogram-like plot with amount of regions (y) per frequency (x). Again the random distribution and the empirical distribution do not differ. **(C, D)** Dependency between Abeta-PET-derived time constant (x) and the probability of dominant theta rhythm (y) across all simulations. **(C)** empirical distribution, **(D)** random distribution. Again we can observe the models criticalities at about 18ms and 24ms, where the theta probability increases and alpha rhythms disappear. Moreover, we can see a shift of the sweet spot for low time constants by the ‘shuffling’: meanwhile for $\tau_i$ =14ms in the empirical distribution theta appears in 70% of the simulations, in the corresponding simulations with random spatial distribution it appears only in 45% of the simulations. This means that the AD-specific spatial pattern leads to specific slowing for regions with very low Abeta in comparison to the random distribution.

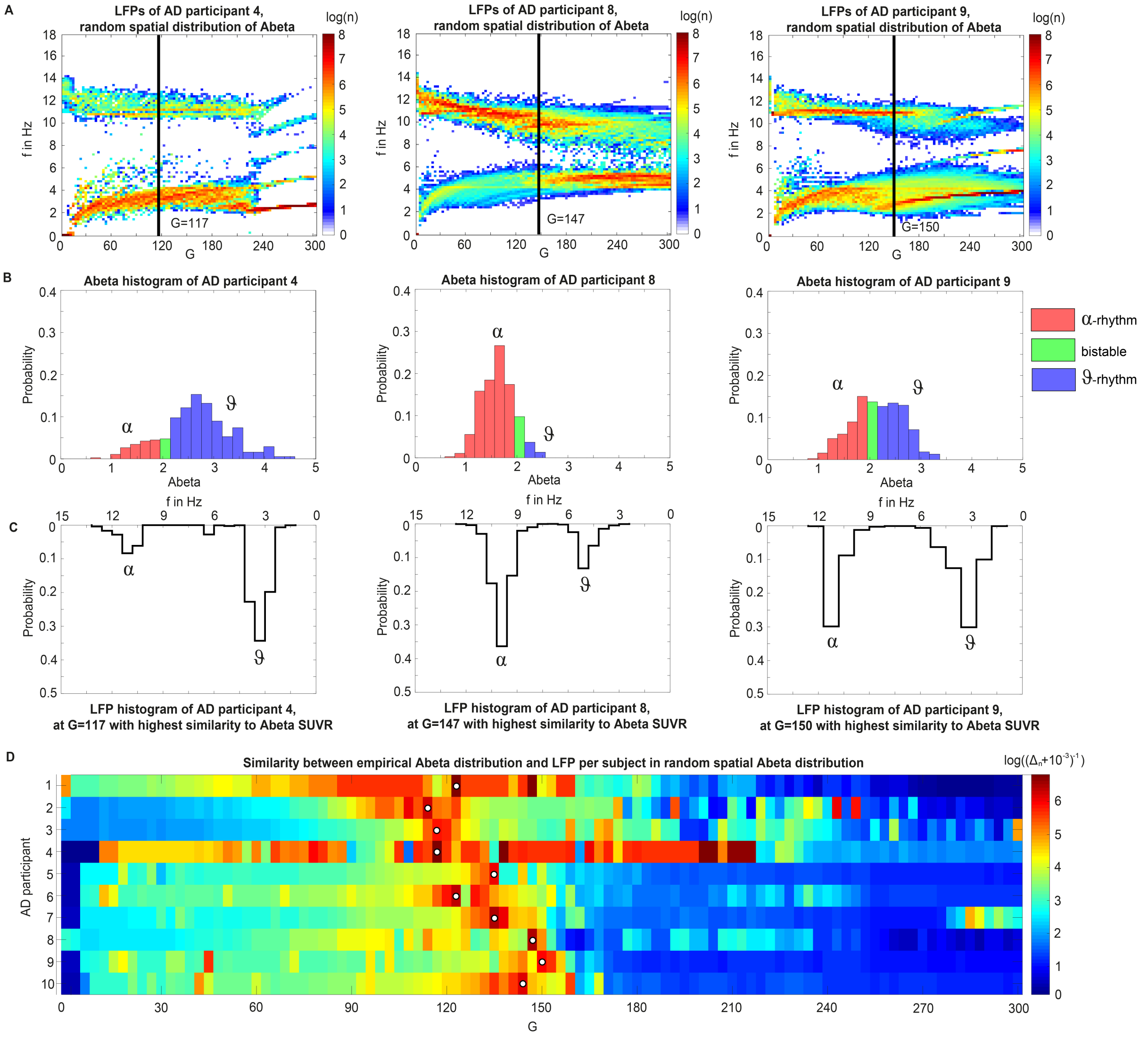


**Supplementary Figure 6. Number of regions in different regimes determine overall spectral properties of simulated EEG.** Surrogate Abeta PET SUVR distributions reveals the role of region burden for the emergence different dynamical regimes. **(A)** shows for three exemplary AD participants (4, 8 and 9) the dominating frequencies (y) of each region depending on global coupling factor G (x axis). Shown is the sum of 10 random spatial distributions of the empirical Abeta . The corresponding histograms with the normalized amount of regions with various Abeta can be seen in **(B)**: classified by their inhibitory time constant to the alpha, theta or bistable regime (**Figure 6**). Below in **(C)** is the frequency distribution shown for a specific value of G, where the ratio of theta rhythm corresponds best to the ratio of regions in theta regime, indicated by a black line in **(A)**. This similarity is demonstrated in **(D)**: the optimal range (white dot) is very similar for all 10 AD subjects at 100<G<150. Similarity is quantified here as the logarithm of the reciprocal of absolute difference between the ratios of theta to alpha and theta together, meaning that high values show high similarity. Note that subject 4 has three additional optima beside the presented one at G = 117: 138, 201, and 204. Beside this optimal correspondence of frequencies and Abeta

SUVRs, we can see a dependency across large parts of the G spectrum: for participant 4, the high amount of theta regime regions lead to a dominant theta cluster in the full G spectrum and synchronizes to a theta/delta rhythm for higher Gs, independent of the spatial distributions. Vice versa, participant 8 has a high amount of alpha regime regions and shows therefore a dominant alpha cluster until G=150 – afterwards the bistable and theta regions reach the theta limit cycle (see **Figure 6**) and propagate slower rhythms. Participant 9 has an equal distribution of alpha and theta regime regions and many bistable regions, which leads to alpha and theta clusters of the same intensity until G=150, while again afterwards the slower rhythms are dominating. The reason for the dominance of slower rhythms for higher G values can also be found in the bifurcation diagrams of **Figure 6**: while in theta regime **(Figure 6I)** for lower Gs the system is in a stable focus and for higher Gs get to the theta limit cycle, the alpha regime **(Figure 6A)** vice versa starts in alpha limit cycle and ends for higher Gs in a stable focus. Since in the absence of noise the system produces no oscillations in stable focuses, for higher Gs the alpha regions with no intrinsic oscillations can easily be synchronized to the propagated theta signal of neighbored regions. All this indicates that TVB weights the distribution of Abeta and transforms it to a complex slowing phenomenon.

### TABLES.

**Supplementary Table 1**. MPRAGE metadata.

| ID | Model | TE [ms] | TR [s] | MatrixSize | VoxelSize [mm] |
|---|---|---|---|---|---|
| 023_S_1190 | TrioTim | 2.98 | 2.3 | (176, 240, 256) | (1.0, 1.0, 1.0) |
| 002_S_1280 | Prisma_fit | 2.95 | 2.3 | (176, 240, 256) | (1.2000046, 1.0546875, 1.0546875) |
| 011_S_4547 | Prisma_fit | 2.98 | 2.3 | (208, 240, 256) | (1.0, 1.0, 1.0) |
| 168_S_6142 | Prisma_fit | 2.98 | 2.3 | (208, 240, 256) | (0.9999948, 1.0, 1.0) |
| 002_S_6103 | Prisma_fit | 2.98 | 2.3 | (208, 240, 256) | (1.0, 1.0, 1.0) |
| 002_S_4654 | Prisma_fit | 2.95 | 2.3 | (176, 240, 256) | (1.199997, 1.0546875, 1.0546875) |
| 022_S_5004 | TrioTim | 2.98 | 2.3 | (176, 240, 256) | (1.0, 1.0, 1.0) |
| 003_S_6067 | Prisma | 2.98 | 2.3 | (208, 240, 256) | (1.0, 1.0, 1.0) |
| 002_S_4229 | Prisma_fit | 2.98 | 2.3 | (208, 240, 256) | (1.0, 1.0, 1.0) |
| 012_S_6073 | Prisma | 2.98 | 2.3 | (208, 240, 256) | (1.0, 1.0, 1.0) |
| 002_S_1261 | Prisma_fit | 2.95 | 2.3 | (176, 240, 256) | (1.2000046, 1.0546875, 1.0546875) |
| 002_S_6009 | Prisma_fit | 2.95 | 2.3 | (176, 240, 256) | (1.2000046, 1.0546875, 1.0546875) |
| 007_S_4488 | Prisma | 2.98 | 2.3 | (208, 240, 256) | (1.0, 1.0, 1.0) |
| 003_S_4288 | Prisma | 2.98 | 2.3 | (208, 240, 256) | (1.0, 1.0, 1.0) |
| 002_S_4213 | Prisma_fit | 2.98 | 2.3 | (208, 240, 256) | (1.0, 1.0, 1.0) |
| 114_S_6039 | Verio | 2.98 | 2.3 | (176, 240, 256) | (1.0, 1.0, 1.0) |
| 036_S_4430 | Skyra | 2.95 | 2.3 | (176, 240, 256) | (1.199997, 1.0546875, 1.0546875) |
| 041_S_4974 | Prisma_fit | 2.95 | 2.3 | (176, 240, 256) | (1.2000046, 1.0546875, 1.0546875) |
| 007_S_4272 | Prisma | 2.98 | 2.3 | (208, 240, 256) | (1.0, 1.0, 1.0) |
| 011_S_4827 | Prisma_fit | 2.98 | 2.3 | (208, 240, 256) | (1.0, 1.0, 1.0) |

| | | | | | |
|---|---|---|---|---|---|
| 002_S_6053 | Prisma_fit | 2.98 | 2.3 | (208, 240, 256) | (1.0, 1.0, 1.0) |
| 003_S_4644 | Prisma | 2.98 | 2.3 | (208, 240, 256) | (1.0, 1.0, 1.0) |
| 002_S_4799 | Prisma_fit | 2.95 | 2.3 | (176, 240, 256) | (1.2000046, 1.0546875, 1.0546875) |
| 002_S_0413 | Prisma_fit | 2.95 | 2.3 | (176, 240, 256) | (1.2000046, 1.0546875, 1.0546875) |
| 114_S_0416 | Verio | 2.98 | 2.3 | (176, 240, 256) | (1.0, 1.0, 1.0) |
| 002_S_5178 | Prisma_fit | 2.95 | 2.3 | (176, 240, 256) | (1.199997, 1.0546875, 1.0546875) |
| 002_S_6030 | Prisma_fit | 2.95 | 2.3 | (176, 240, 256) | (1.2000046, 1.0546875, 1.0546875) |
| 003_S_1122 | Prisma | 2.98 | 2.3 | (208, 240, 256) | (1.0000056, 1.0, 1.0) |
| 011_S_4893 | Prisma_fit | 2.98 | 2.3 | (208, 240, 256) | (1.0, 1.0, 1.0) |
| 002_S_1155 | Prisma_fit | 2.95 | 2.3 | (176, 240, 256) | (1.2000046, 1.0546875, 1.0546875) |
| 036_S_4715 | Skyra | 2.95 | 2.3 | (176, 240, 256) | (1.2000046, 1.0546875, 1.0546875) |
| 007_S_4387 | Prisma | 2.98 | 2.3 | (208, 240, 256) | (1.0, 1.0, 1.0) |
| 007_S_4620 | Prisma | 2.98 | 2.3 | (208, 240, 256) | (1.0, 1.0, 1.0) |

**Supplementary Table 2**. FLAIR metadata.

| ID | Model | TE [ms] | TR [s] | MatrixSize | VoxelSize [mm] |
|---|---|---|---|---|---|
| 023_S_1190 | TrioTim | 443 | 4.8 | (160, 256, 256) | (1.2000046, 1.0, 1.0) |
| 002_S_1280 | Prisma_fit | 441 | 4.8 | (160, 256, 256) | (1.2000046, 1.0, 1.0) |
| 011_S_4547 | Prisma_fit | 441 | 4.8 | (160, 256, 256) | (1.2000046, 1.0, 1.0) |
| 168_S_6142 | Prisma_fit | 441 | 4.8 | (160, 256, 256) | (1.2000002, 1.0, 1.0) |
| 002_S_6103 | Prisma_fit | 441 | 4.8 | (160, 256, 256) | (1.199997, 1.0, 1.0) |
| 002_S_4654 | Prisma_fit | 441 | 4.8 | (160, 256, 256) | (1.199997, 1.0, 1.0) |
| 022_S_5004 | TrioTim | 439 | 4.8 | (160, 256, 256) | (1.199997, 1.0, 1.0) |
| 003_S_6067 | Prisma | 441 | 4.8 | (160, 256, 256) | (1.2000046, 1.0, 1.0) |
| 002_S_4229 | Prisma_fit | 441 | 4.8 | (160, 256, 256) | (1.199997, 1.0, 1.0) |
| 012_S_6073 | Prisma | 441 | 4.8 | (160, 256, 256) | (1.2000046, 1.0, 1.0) |
| 002_S_1261 | Prisma_fit | 441 | 4.8 | (160, 256, 256) | (1.2000046, 1.0, 1.0) |
| 002_S_6009 | Prisma_fit | 441 | 4.8 | (160, 256, 256) | (1.2000046, 1.0, 1.0) |
| 007_S_4488 | Prisma | 441 | 4.8 | (160, 256, 256) | (1.2000046, 1.0, 1.0) |
| 003_S_4288 | Prisma | 441 | 4.8 | (160, 256, 256) | (1.199997, 1.0, 1.0) |
| 002_S_4213 | Prisma_fit | 441 | 4.8 | (160, 256, 256) | (1.199997, 1.0, 1.0) |
| 114_S_6039 | Verio | 343 | 4.8 | (160, 256, 256) | (1.0, 1.0, 1.0) |
| 036_S_4430 | Skyra | 441 | 4.8 | (160, 256, 256) | (1.2000046, 1.0, 1.0) |
| 041_S_4974 | Prisma_fit | 441 | 4.8 | (160, 256, 256) | (1.199997, 1.0, 1.0) |
| 007_S_4272 | Prisma | 441 | 4.8 | (160, 256, 256) | (1.199997, 1.0, 1.0) |
| 011_S_4827 | Prisma_fit | 441 | 4.8 | (160, 256, 256) | (1.199997, 1.0, 1.0) |

| | | | | | |
|---|---|---|---|---|---|
| 002_S_6053 | Prisma_fit | 441 | 4.8 | (160, 256, 256) | (1.199997, 1.0, 1.0) |
| 003_S_4644 | Prisma | 441 | 4.8 | (160, 256, 256) | (1.199997, 1.0, 1.0) |
| 002_S_4799 | Prisma_fit | 441 | 4.8 | (160, 256, 256) | (1.2000046, 1.0, 1.0) |
| 002_S_0413 | Prisma_fit | 441 | 4.8 | (160, 256, 256) | (1.2000046, 1.0, 1.0) |
| 114_S_0416 | Verio | 343 | 4.8 | (160, 256, 256) | (1.0, 1.0, 1.0) |
| 002_S_5178 | Prisma_fit | 441 | 4.8 | (160, 256, 256) | (1.199997, 1.0, 1.0) |
| 002_S_6030 | Prisma_fit | 441 | 4.8 | (160, 256, 256) | (1.2000046, 1.0, 1.0) |
| 003_S_1122 | Prisma | 441 | 4.8 | (160, 256, 256) | (1.2000005, 1.0, 1.0) |
| 011_S_4893 | Prisma_fit | 441 | 4.8 | (160, 256, 256) | (1.2000046, 1.0, 1.0) |
| 002_S_1155 | Prisma_fit | 441 | 4.8 | (160, 256, 256) | (1.2000046, 1.0, 1.0) |
| 036_S_4715 | Skyra | 441 | 4.8 | (160, 256, 256) | (1.2000046, 1.0, 1.0) |
| 007_S_4387 | Prisma | 441 | 4.8 | (160, 256, 256) | (1.2000046, 1.0, 1.0) |
| 007_S_4620 | Prisma | 441 | 4.8 | (160, 256, 256) | (1.2000046, 1.0, 1.0) |

**Supplementary Table 3**. DTI metadata (only for HC participants to average the SC template)

| ID | Model | Institute | TE [ms] | TR [s] | MatrixSize | VoxelSize [mm, mm, mm, s] | n_Bvecs | Bvals |
|---|---|---|---|---|---|---|---|---|
| 002_S_1280 | Prisma_fit | OHSU_AIRC | 56 | 7.2 | (116, 116, 80, 55) | (2.0, 2.0, 2.0, 7.2) | 49 | [ 0. 1000.] |
| 002_S_6103 | Prisma_fit | OHSU_AIRC | 56 | 7.2 | (116, 116, 80, 55) | (2.0, 2.0, 2.0, 7.2) | 49 | [ 0. 1000.] |
| 003_S_6067 | Prisma | USCINI | 56 | 7.2 | (116, 116, 80, 55) | (2.0, 2.0, 2.0, 7.2) | 49 | [ 0. 1000.] |
| 002_S_6009 | Prisma_fit | OHSU_AIRC | 56 | 7.2 | (116, 116, 80, 55) | (2.0, 2.0, 2.0, 7.2) | 49 | [ 0. 1000.] |
| 007_S_4488 | Prisma | MAYO_CLINIC_MRI_58 | 71 | 3.4 | (116, 116, 81, 127) | (2.0, 2.0, 2.0, 3.4) | 115 | [ 0. 500. 1000. 2000.] |
| 003_S_4288 | Prisma | USC_Stevens_Hall_Institute | 56 | 7.2 | (116, 116, 80, 55) | (2.0, 2.0, 2.0, 7.2) | 49 | [ 0. 1000.] |
| 002_S_4213 | Prisma_fit | OHSU_AIRC | 56 | 7.2 | (116, 116, 80, 55) | (2.5172415, 2.5172415, 2.0, 7.2) | 49 | [ 0. 1000.] |
| 002_S_6053 | Prisma_fit | OHSU_AIRC | 56 | 7.2 | (116, 116, 80, 55) | (2.0, 2.0, 2.0, 7.2) | 49 | [ 0. 1000.] |
| 003_S_4644 | Prisma | USCINI | 56 | 7.2 | (116, 116, 80, 55) | (2.0, 2.0, 2.0, 7.2) | 49 | [ 0. 1000.] |
| 002_S_4799 | Prisma_fit | OHSU_AIRC | 56 | 7.2 | (116, 116, 80, 55) | (2.0, 2.0, 2.0, 7.2) | 49 | [ 0. 1000.] |
| 002_S_0413 | Prisma_fit | OHSU_AIRC | 56 | 7.2 | (116, 116, 80, 55) | (2.0, 2.0, 2.0, 7.2) | 49 | [ 0. 1000.] |
| 002_S_5178 | Prisma_fit | OHSU_AIRC | 56 | 7.2 | (116, 116, 80, 55) | (2.0, 2.0, 2.0, 7.2) | 49 | [ 0. 1000.] |
| 002_S_6030 | Prisma_fit | OHSU_AIRC | 56 | 7.2 | (116, 116, 80, 55) | (2.0, 2.0, 2.0, 7.2) | 49 | [ 0. 1000.] |
| 007_S_4387 | Prisma | MAYO_CLINIC_MRI_58 | 71 | 3.4 | (116, 116, 81, 127) | (2.0, 2.0, 2.0, 3.4) | 115 | [ 0. 500. 1000. 2000.] |
| 007_S_4620 | Prisma | MAYO_CLINIC_MRI_58 | 71 | 3.4 | (116, 116, 81, 127) | (2.0, 2.0, 2.0, 3.4) | 115 | [ 0. 500. 1000. 2000.] |

**Supplementary Table 4.** AV-45 PET (Abeta) metadata.

| ID | Scanner | Model | MatrixSize | VoxelSize [mm] |
|---|---|---|---|---|

| | | | | |
|---|---|---|---|---|
| 023_S_1190 | Siemens | Biograph6_TruePoint | (160, 160, 96) | (1.5, 1.5, 1.5) |
| 002_S_1280 | Philips | GEMINI_TF_TOF_16 | (160, 160, 96) | (1.5, 1.5, 1.5) |
| 011_S_4547 | Siemens | Biograph40_TruePoint | (160, 160, 96) | (1.5, 1.5, 1.5) |
| 168_S_6142 | GE | Discovery_STE | (160, 160, 96) | (1.5, 1.5, 1.5) |
| 002_S_6103 | Philips | GEMINI_TF_TOF_16 | (160, 160, 96) | (1.5, 1.5, 1.5) |
| 002_S_4654 | Philips | GEMINI_TF_TOF_16 | (160, 160, 96) | (1.5, 1.5, 1.5) |
| 022_S_5004 | Philips | Ingenuity_TF_PET_CT | (160, 160, 96) | (1.5, 1.5, 1.5) |
| 003_S_6067 | Siemens | Biograph64_TruePoint | (160, 160, 96) | (1.5, 1.5, 1.5) |
| 002_S_4229 | Philips | GEMINI_TF_TOF_16 | (160, 160, 96) | (1.5, 1.5, 1.5) |
| 012_S_6073 | GE | Discovery_710 | (160, 160, 96) | (1.5, 1.5, 1.5) |
| 002_S_1261 | Philips | GEMINI_TF_TOF_16 | (160, 160, 96) | (1.5, 1.5, 1.5) |
| 002_S_6009 | Philips | GEMINI_TF_TOF_16 | (160, 160, 96) | (1.5, 1.5, 1.5) |
| 007_S_4488 | GE | Discovery_690 | (160, 160, 96) | (1.5, 1.5, 1.5) |
| 003_S_4288 | Siemens | Biograph64_TruePoint | (160, 160, 96) | (1.5, 1.5, 1.5) |
| 002_S_4213 | Philips | GEMINI_TF_TOF_16 | (160, 160, 96) | (1.5, 1.5, 1.5) |
| 114_S_6039 | Philips | GEMINI_TF_TOF_64 | (160, 160, 96) | (1.5, 1.5, 1.5) |
| 036_S_4430 | Siemens | Biograph40_TruePoint | (160, 160, 96) | (1.5, 1.5, 1.5) |
| 041_S_4974 | Siemens | HR+ | (160, 160, 96) | (1.5, 1.5, 1.5) |
| 007_S_4272 | GE | Discovery_690 | (160, 160, 96) | (1.5, 1.5, 1.5) |
| 011_S_4827 | Siemens | Biograph40_TruePoint | (160, 160, 96) | (1.5, 1.5, 1.5) |
| 002_S_6053 | Philips | GEMINI_TF_TOF_16 | (160, 160, 96) | (1.5, 1.5, 1.5) |
| 003_S_4644 | Siemens | Biograph64_TruePoint | (160, 160, 96) | (1.5, 1.5, 1.5) |
| 002_S_4799 | Philips | GEMINI_TF_TOF_16 | (160, 160, 96) | (1.5, 1.5, 1.5) |
| 002_S_0413 | Philips | GEMINI_TF_TOF_16 | (160, 160, 96) | (1.5, 1.5, 1.5) |
| 114_S_0416 | Philips | GEMINI_TF_TOF_64 | (160, 160, 96) | (1.5, 1.5, 1.5) |
| 002_S_5178 | Philips | GEMINI_TF_TOF_16 | (160, 160, 96) | (1.5, 1.5, 1.5) |
| 002_S_6030 | Philips | GEMINI_TF_TOF_16 | (160, 160, 96) | (1.5, 1.5, 1.5) |
| 003_S_1122 | Siemens | Biograph64_TruePoint | (160, 160, 96) | (1.5, 1.5, 1.5) |
| 011_S_4893 | Siemens | Biograph40_TruePoint | (160, 160, 96) | (1.5, 1.5, 1.5) |
| 002_S_1155 | Philips | GEMINI_TF_TOF_16 | (160, 160, 96) | (1.5, 1.5, 1.5) |
| 036_S_4715 | Siemens | Biograph40_TruePoint | (160, 160, 96) | (1.5, 1.5, 1.5) |
| 007_S_4387 | GE | Discovery_690 | (160, 160, 96) | (1.5, 1.5, 1.5) |
| 007_S_4620 | GE | Discovery_690 | (160, 160, 96) | (1.5, 1.5, 1.5) |

**Supplementary Table 5.** AV-14-51 PET (Tau) metadata

| ID | Scanner | Model | MatrixSize | VoxelSize [mm] |
|---|---|---|---|---|
| 023_S_1190 | Siemens | Biograph6_TruePoint | (160, 160, 96) | (1.5, 1.5, 1.5) |
| 002_S_1280 | Philips | GEMINI_TF_TOF_16 | (160, 160, 96) | (1.5, 1.5, 1.5) |
| 011_S_4547 | Siemens | Biograph40_TruePoint | (160, 160, 96) | (1.5, 1.5, 1.5) |
| 168_S_6142 | GE | Discovery_STE | (160, 160, 96) | (1.5, 1.5, 1.5) |
| 002_S_6103 | Philips | GEMINI_TF_TOF_16 | (160, 160, 96) | (1.5, 1.5, 1.5) |
| 002_S_4654 | Philips | GEMINI_TF_TOF_16 | (160, 160, 96) | (1.5, 1.5, 1.5) |
| 022_S_5004 | Philips | Ingenuity_TF_PET_CT | (160, 160, 96) | (1.5, 1.5, 1.5) |
| 003_S_6067 | Siemens | Biograph64_TruePoint | (160, 160, 96) | (1.5, 1.5, 1.5) |
| 002_S_4229 | Philips | GEMINI_TF_TOF_16 | (160, 160, 96) | (1.5, 1.5, 1.5) |
| 012_S_6073 | GE | Discovery_710 | (160, 160, 96) | (1.5, 1.5, 1.5) |
| 002_S_1261 | Philips | GEMINI_TF_TOF_16 | (160, 160, 96) | (1.5, 1.5, 1.5) |
| 002_S_6009 | Philips | GEMINI_TF_TOF_16 | (160, 160, 96) | (1.5, 1.5, 1.5) |
| 007_S_4488 | GE | Discovery_690 | (160, 160, 96) | (1.5, 1.5, 1.5) |
| 003_S_4288 | Siemens | Biograph64_TruePoint | (160, 160, 96) | (1.5, 1.5, 1.5) |
| 002_S_4213 | Philips | GEMINI_TF_TOF_16 | (160, 160, 96) | (1.5, 1.5, 1.5) |
| 114_S_6039 | Philips | GEMINI_TF_TOF_64 | (160, 160, 96) | (1.5, 1.5, 1.5) |
| 036_S_4430 | Siemens | Biograph40_TruePoint | (160, 160, 96) | (1.5, 1.5, 1.5) |
| 041_S_4974 | Siemens | HR+ | (160, 160, 96) | (1.5, 1.5, 1.5) |
| 007_S_4272 | GE | Discovery_690 | (160, 160, 96) | (1.5, 1.5, 1.5) |
| 011_S_4827 | Siemens | Biograph40_TruePoint | (160, 160, 96) | (1.5, 1.5, 1.5) |
| 002_S_6053 | Philips | GEMINI_TF_TOF_16 | (160, 160, 96) | (1.5, 1.5, 1.5) |
| 003_S_4644 | Siemens | Biograph64_TruePoint | (160, 160, 96) | (1.5, 1.5, 1.5) |
| 002_S_4799 | Philips | GEMINI_TF_TOF_16 | (160, 160, 96) | (1.5, 1.5, 1.5) |
| 002_S_0413 | Philips | GEMINI_TF_TOF_16 | (160, 160, 96) | (1.5, 1.5, 1.5) |
| 114_S_0416 | Philips | GEMINI_TF_TOF_64 | (160, 160, 96) | (1.5, 1.5, 1.5) |
| 002_S_5178 | Philips | GEMINI_TF_TOF_16 | (160, 160, 96) | (1.5, 1.5, 1.5) |
| 002_S_6030 | Philips | GEMINI_TF_TOF_16 | (160, 160, 96) | (1.5, 1.5, 1.5) |
| 003_S_1122 | Siemens | Biograph64_TruePoint | (160, 160, 96) | (1.5, 1.5, 1.5) |
| 011_S_4893 | Siemens | Biograph40_TruePoint | (160, 160, 96) | (1.5, 1.5, 1.5) |
| 002_S_1155 | Philips | GEMINI_TF_TOF_16 | (160, 160, 96) | (1.5, 1.5, 1.5) |
| 036_S_4715 | Siemens | Biograph40_TruePoint | (160, 160, 96) | (1.5, 1.5, 1.5) |
| 007_S_4387 | GE | Discovery_690 | (160, 160, 96) | (1.5, 1.5, 1.5) |
| 007_S_4620 | GE | Discovery_690 | (160, 160, 96) | (1.5, 1.5, 1.5) |

**Supplementary Table 6.** Dates of Imaging and MMSE.

| ID | MMSE date | MPRAGE date | FLAIR date | DTI date | AV-45 PET date | AV-1451 PET date |
|---|---|---|---|---|---|---|
| 023_S_1190 | 17/11/13 | 17/10/23 | 17/10/23 | | 17/10/25 | 17/11/08 |
| 002_S_1280 | 18/3/7 | 17/3/13 | 17/3/13 | 17/3/13 | 17/3/02 | 18/3/5 |
| 011_S_4547 | 17/8/18 | 17/8/18 | 17/8/18 | | 17/8/30 | 17/8/24 |
| 168_S_6142 | 17/12/5 | 17/12/18 | 17/12/18 | | 18/1/17 | 18/1/3 |
| 002_S_6103 | 17/10/25 | 17/11/20 | 17/11/20 | 17/11/20 | 17/11/21 | 18/1/17 |
| 002_S_4654 | 18/5/15 | 17/5/3 | 17/5/3 | | 17/5/2 | 18/5/22 |
| 022_S_5004 | 18/6/29 | 18/3/14 | 17/3/21 | | 17/3/21 | 17/4/5 |
| 003_S_6067 | 17/12/4 | 17/8/18 | 17/8/18 | 17/8/18 | 17/10/13 | 17/10/18 |
| 002_S_4229 | 18/5/14 | 17/9/20 | 17/9/20 | | 17/9/20 | 17/10/3 |
| 012_S_6073 | 17/9/18 | 17/9/22 | 17/9/22 | | 17/10/12 | 17/10/11 |
| 002_S_1261 | 18/3/8 | 17/3/15 | 17/3/15 | | 17/3/14 | 17/3/15 |
| 002_S_6009 | 17/4/1 | 17/4/17 | 17/4/17 | 17/4/17 | 17/5/16 | 17/5/15 |
| 007_S_4488 | 18/6/11 | 17/9/12 | 17/9/12 | 17/9/12 | 17/9/22 | 17/9/13 |
| 003_S_4288 | 17/10/2 | 17/10/3 | 17/10/3 | 17/10/3 | 17/10/3 | 18/2/22 |
| 002_S_4213 | 17/8/16 | 17/8/14 | 17/8/14 | 17/8/14 | 17/8/14 | 17/8/17 |
| 114_S_6039 | 17/8/10 | 17/7/21 | 17/7/21 | | 17/8/24 | 17/10/4 |
| 036_S_4430 | 17/11/15 | 17/11/07 | 17/11/07 | | 17/11/15 | 17/11/21 |
| 041_S_4974 | 17/10/30 | 17/10/5 | 17/10/5 | | 17/8/24 | 17/10/12 |
| 007_S_4272 | 18/1/18 | 18/1/16 | 18/1/16 | | 17/12/19 | 18/1/17 |
| 011_S_4827 | 17/8/24 | 17/8/31 | 17/8/31 | | 17/8/28 | 17/9/7 |
| 002_S_6053 | 17/7/21 | 17/7/18 | 17/7/18 | 17/7/18 | 17/8/23 | 17/8/24 |
| 003_S_4644 | 17/6/26 | 17/6/21 | 17/6/21 | 17/6/21 | 18/2/28 | 18/4/17 |
| 002_S_4799 | 18/6/7 | 17/5/22 | 17/5/22 | 17/5/22 | 17/5/18 | 18/6/13 |
| 002_S_0413 | 17/6/16 | 17/6/21 | 17/6/21 | 17/6/21 | 17/6/15 | 17/6/21 |
| 114_S_0416 | 18/7/24 | 17/10/24 | 17/10/24 | | 17/10/24 | 17/11/21 |
| 002_S_5178 | 17/6/6 | 17/5/31 | 17/5/31 | 17/5/31 | 17/6/5 | 17/5/31 |
| 002_S_6030 | 17/6/9 | 17/6/15 | 17/6/15 | 17/6/15 | 17/7/25 | 17/7/24 |
| 003_S_1122 | 18/7/25 | 17/5/18 | 17/5/18 | | 17/8/8 | 17/8/10 |
| 011_S_4893 | 18/7/17 | 17/11/8 | 17/11/8 | | 17/11/1 | 17/11/7 |
| 002_S_1155 | 18/5/9 | 17/4/24 | 17/4/24 | | 17/4/20 | 17/4/24 |
| 036_S_4715 | 17/10/13 | 17/10/10 | 17/10/10 | | 17/10/10 | 17/10/12 |
| 007_S_4387 | 17/10/31 | 17/11/1 | 17/11/1 | 17/11/1 | 17/10/24 | 17/11/29 |
| 007_S_4620 | 17/12/12 | 17/12/05 | 17/12/05 | 17/12/05 | 17/12/06 | 17/12/14 |

**Supplementary Table 7.** Participants IDs used in this study and corresponding ADNI official ID.

| | ID | ADNI ID |
|---|---|---|
| AD | 1 | 023_S_1190 |
| | 2 | 011_S_4547 |
| | 3 | 168_S_6142 |
| | 4 | 114_S_6039 |
| | 5 | 036_S_4430 |
| | 6 | 041_S_4974 |
| | 7 | 011_S_4827 |
| | 8 | 114_S_0416 |
| | 9 | 011_S_4893 |
| | 10 | 036_S_4715 |
| HC | 11 | 002_S_1280 |
| | 12 | 002_S_6103 |
| | 13 | 003_S_6067 |
| | 14 | 002_S_6009 |
| | 15 | 007_S_4488 |
| | 16 | 003_S_4288 |
| | 17 | 002_S_4213 |
| | 18 | 002_S_6053 |
| | 19 | 003_S_4644 |
| | 20 | 002_S_4799 |
| | 21 | 002_S_0413 |
| | 22 | 002_S_5178 |
| | 23 | 002_S_6030 |
| | 24 | 007_S_4387 |
| | 25 | 007_S_4620 |
| MCI | 26 | 002_S_4654 |
| | 27 | 022_S_5004 |
| | 28 | 002_S_4229 |
| | 29 | 012_S_6073 |
| | 30 | 002_S_1261 |
| | 31 | 007_S_4272 |
| | 32 | 003_S_1122 |
| | 33 | 002_S_1155 |

1. Spiegler A, Kiebel SJ, Atay FM, Knösche TR. Bifurcation analysis of neural mass models: Impact of extrinsic inputs and dendritic time constants. NeuroImage. 2010;52(3):1041-58.
2. Strogatz SH. Nonlinear dynamics and chaos : with applications to physics, biology, chemistry, and engineering: Second edition. Boulder, CO : Westview Press, a member of the Perseus Books Group, [2015]; 2015.
